# Cross-tissue single-cell transcriptomics reveals organizing principles of fibroblasts in health and disease

**DOI:** 10.1101/2021.01.15.426912

**Authors:** Matthew B. Buechler, Rachana N. Pradhan, Aslihan Karabacak Calviello, Soren Muller, Richard Bourgon, Shannon. J. Turley

## Abstract

Fibroblasts are non-hematopoietic structural cells that define the architecture of organs, support the homeostasis of tissue-resident cells and play key roles in fibrosis, cancer, autoimmunity and wound healing. Recent studies have described fibroblast heterogeneity within individual tissues. However, the field lacks a definition of fibroblasts at single-cell resolution across tissues in healthy and diseased organs. Here, we integrated single-cell RNA transcriptomic data from ~150,000 fibroblast cells derived from 16 steady- and 11 perturbed-state mouse organs into fibroblast atlases. These data revealed two universal fibroblast cell subtypes, marked by expression of *Pi16* or *Col15a1*, in all tissues; it also revealed discrete subsets of five specialized fibroblast subtypes in steady-state tissues and three activated fibroblast subtypes in perturbed or diseased tissues. These subsets were transcriptionally shaped by microenvironmental context rather than tissue-type alone. Inference of fibroblast lineage structure from the murine steady-state and perturbed-state fibroblast atlases suggested that specialized and activated subtypes are developmentally related to universal tissue-resident fibroblasts. Analysis of human samples revealed that fibroblast subtypes found in mice are conserved between species, including universal fibroblasts and activated phenotypes associated with pathogenicity in human cancer, fibrosis, arthritis and inflammation. In sum, a cross-species and pan-tissue approach to transcriptomics at single-cell resolution enabled us to define the organizing principles of the fibroblast lineage in health and disease.

## Introduction

Fibroblasts populate all tissues of the body, delineate the topography of organs via the production and remodeling of extracellular matrix proteins (ECMs)^1^ and support tissue-resident cell types including macrophages, T cells, B cells, innate-like lymphocytes, dendritic cells, hematopoietic stem cells and endothelial cells^2,3^. These mesenchymal cells appear to perform both general functions associated with their lineage regardless of context and specialized programs suited to the needs of the tissue to maintain organ homeostasis. Other cell types such as macrophages achieve parallel generalized function and specialization through expression of a lineage-wide core transcriptomic signature accompanied by tissue-specific programming driven by microenvironmental cues^4–6^. How fibroblasts execute functions common to their lineage but also as required by their organ of residence is not clear.

Technological advances, such as single-cell RNA-sequencing (scRNA-seq), have revealed an intriguing degree of intra-tissue fibroblast heterogeneity^2^, though whether these insights apply across tissues is unknown. Understanding fibroblast inter-tissue population structure has clinical relevance as subtypes of fibroblasts drive inflammation and tissue destruction in arthritis^7–10^, promote malignancy in cancer^11–15^ and deposit ECMs that promotes tissue dysfunction in fibrotic indications such as idiopathic pulmonary fibrosis (IPF)^1^. Emerging paradigms within the arthritis^7^ and cancer^12,16^ fields have suggested that two discrete fibroblast subtypes drive aspects of disease: (i) an inflammatory fibroblast that secretes inflammatory factors that recruit immune cells and (ii) a contractile myofibroblast which mediates tissue destruction in arthritis and immune exclusion in solid tumors. Revealing whether fibroblast phenotypes across indications are wholly context specific or in some cases share molecular similarities may help to inform therapeutic approaches that target these cells. In this study, we hypothesized that pantissue fibroblast phenotypes exist but that this lineage achieves context-specific functionality via cell subtypes that develop in response to cues derived from healthy and perturbed local milieus.

## Results

### Cross-tissue transcriptional panorama of fibroblasts in steady-state mouse tissues

To investigate this hypothesis, we first performed bulk RNA-seq and Assay for Transposase-Accessible Chromatin (ATAC)-seq on fibroblasts that were FACS-sorted as previously published^17^ from a range of mouse tissues. These data revealed transcriptional cassettes, regions of open chromatin and transcriptional networks that were driven by tissue-type (Extended Data Figs. 1–2, Supplementary Tables 1-2), similar to recent reports^17,18^. Bulk sequencing modalities cannot determine whether derived signatures are reflective of a single, homogenous population within a tissue or an average of heterogeneous populations. To resolve this, we utilized scRNA-seq. We collected a compendium of scRNA-seq datasets enriched for non-hematopoietic cells from our laboratory and public repositories that were processed using the 10X Genomics platform. We ensured consistency in detected genes and mitochondrial read counts and then removed hematopoietic, endothelial, mesothelial and mural (pericyte and smooth muscle) cells and corrected for cross-laboratory batch effects with the *Harmony*^19^ algorithm, resulting in a fibroblast-specific single-cell map (Extended Data Fig. 3a-c). This compilation, which we termed the steady-state fibroblast atlas, was composed of 21 datasets across 16 different unperturbed tissues^12,20–38^ and contained 86,379 cells (Fig. 1a-b and Supplementary Table 3). An interactive data browser for the steady-state fibroblast atlas and other content from this manuscript is publicly available (for details see ‘Data availability’). To demonstrate concordance between our bulk fibroblast RNA-seq data and the single-cell atlas, we computed bulk tissue-specific fibroblast signatures and scored the average single-cell expression profiles tissue-by-tissue. Profiles were highly concordant, indicating our analytical approach to the public single-cell datasets did not introduce technical bias (Extended Data Fig. 3d). Moreover, it suggested that single-cell data could both explain the bulk sequencing patterns we observed and provide substantially more clarity on the nature of fibroblast subtypes within and across tissues.

**Fig 1.**
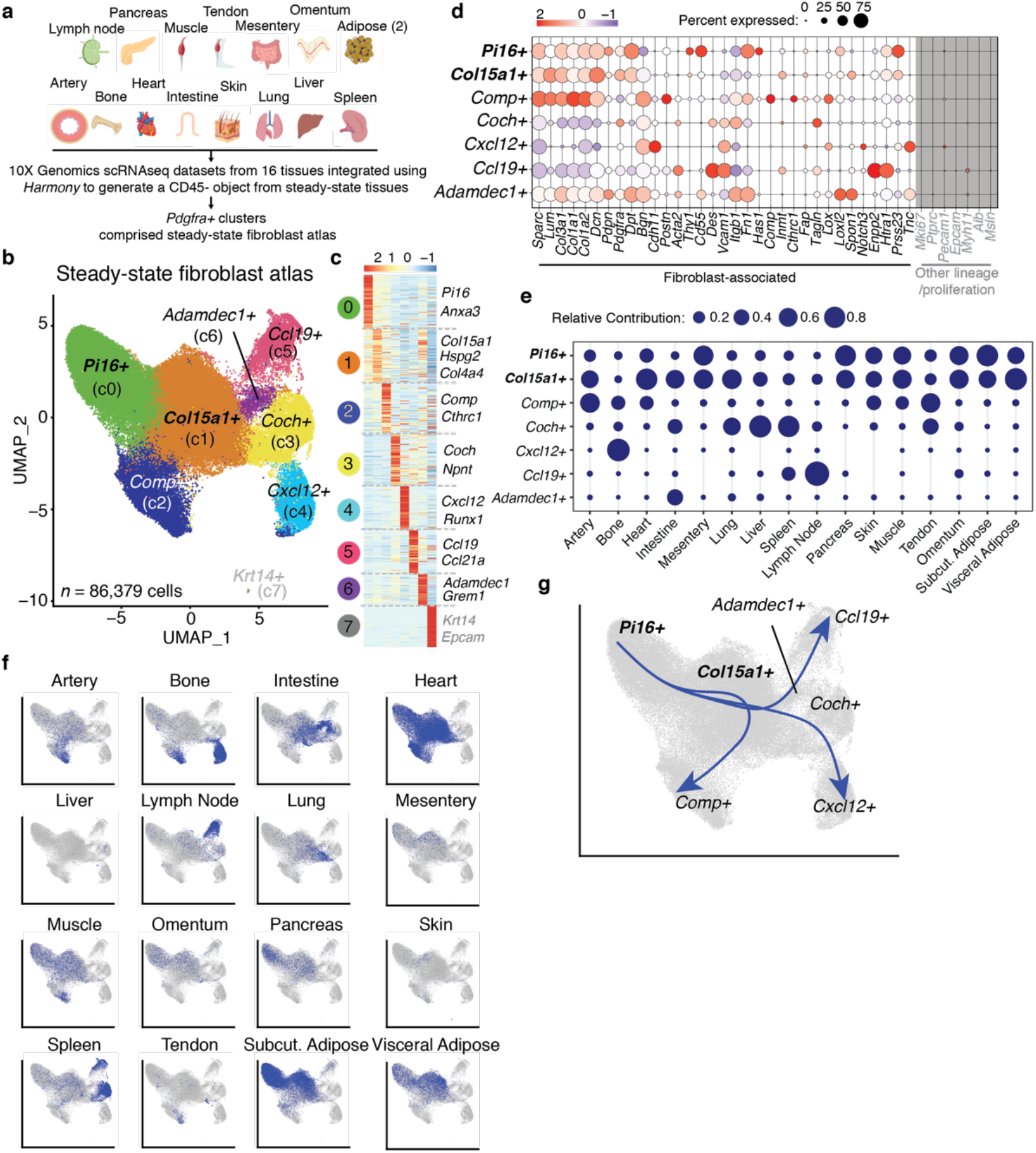
Cross-tissue transcriptional panorama of fibroblasts in resting mouse tissues. **a**. Schematic of datasets compiled for steady-state fibroblast atlas from 16 mouse tissues: artery, bone and bone marrow, heart, intestine, skin, lung, liver, spleen, lymph node, pancreas, muscle, tendon, mesentery, omentum and visceral and subcutaneous adipose. **b**. UMAP embedding of 86,379 single cells in steady-state fibroblast atlas. Eight clusters identified through graph-based clustering are indicated by color. **c**. Heat map of the relative average expression of the most strongly enriched genes for each cluster identified by log-fold change of cells in one cluster compared to all other cells (z-score per row). **d**. Fibroblast- and other lineage-associated genes (in grey). The size of circles denotes the percentage of cells from each cluster, color encodes the average expression across all cells within a cluster. The color scale shows the expression level based on row z-score. **e**. Relative abundance of each tissue in steady-state UMAP clusters. The size of bubbles indicates the contribution of cells from each tissue to a cluster, columns without a bubble indicate lack of contribution (number of cells = 0) of that tissue to the corresponding cluster. Graph to be read column-wise. **f**. UMAPs highlighting distribution of cells from individual tissues in the steady-state fibroblast atlas. **g**. Pseudotime(s) visualized using principal curves representing three trajectories of fibroblast differentiation across steady-state fibroblast object. *Pi16+* cluster set as root fibroblast state.

In the steady-state fibroblast atlas, seven clusters of fibroblasts and one cluster of epithelial contaminants (*Krt14+*) were identified based upon differential gene expression analysis at a clustering resolution selected to unify similar cell states (Fig. 1c). This approach identified greater than 140 marker genes for each cluster and we chose to annotate clusters based on the dominant cluster-specific gene: *Pi16+, Col15a1+, Comp+, Coch+, Cxcl12+, Ccl19+* and *Adamdec1+* (Supplementary Table 4 and Extended Data Fig. 3e). We noted that these clusters were transcriptionally distinct based upon known fibroblast-associated genes, revealing intriguing heterogeneity within the fibroblast lineage (Fig. 3d). All tissues contributed to the *Pi16+* and *Col15a1+* clusters, suggesting that these clusters are universal fibroblasts (Fig. 1e-f). Consistent with this characterization and the hypothesized universal presence across tissues, the genes defining these two clusters strongly differentiated fibroblasts from mesothelial cells in bulk RNAseq data from six different tissue sources (Extended Data Fig. 4). Some other clusters included cells from multiple but not all tissues, whereas two clusters, *Adamdec1+* (intestine) and *Cxcl12+* (bone), derived greater than 90% of cells from just one tissue (Fig. 1e-f). The more limited sets of tissues contributing to the *Ccl19+, Coch+, Cxcl12+, Comp+* and *Adamdec1+*clusters relative to universal fibroblast clusters suggested these may represent more specialized cell states.

To achieve a deeper understanding of the fibroblast clusters identified in the fibroblast steady-state atlas, we first performed gene set enrichment analysis of all the clusters (Extended Data Fig. 5a) and then compared the *Ccl19+, Coch+, Cxcl12+, Adamdec1+* and *Comp+* clusters to the universal *Pi16+* and *Col15a1+* clusters (Extended Data Fig. 5b-f, Supplementary Table 4). This approach suggested that the *Ccl19+* cluster was composed of fibroblastic reticular cells, which regulate immune cell localization and function in secondary lymphoid organs^3^. Cells within the *Coch+* cluster corresponded to red pulp fibroblasts and alveolar fibroblasts, as determined by higher expression of *Wt1, Csf1* (red pulp fibroblasts) and *Npnt* and *Ces1d* (alveolar fibroblasts)^34,39–41^ relative to universal fibroblasts. This is consistent with observations showing that *Wt1*-expressing red pulp fibroblasts maintain red pulp macrophage homeostasis via provision of *Csf1*^39^, though it is unclear if these mechanisms apply in the lung. The *Cxcl12+* cluster was a mix of *Lepr^+^* mesenchymal stromal cells which express *Lepr* and *Adipoq* and mesenchymal-derived *Emb+, Sp7+* osteolineage cells^21,42–44^, two cell types that are known to regulate hematopoietic niches in the bone microenvironment. The *Adamdec1+* cluster consisted of ECM-depositing cells that are known to produce key niche factors for intestinal stem cells^22,45^. The *Comp+* cluster represented the ‘fibroblast 2’ cell state in the artery, a fibroblast cell population that has previously been described as lacking expression of *Pi16, Clec3b, Dpep1,* consistent with our data, but does not have a functional annotation^20^. The specialization of these clusters was reflected in differential enrichment of core signaling pathways, including NFkB and TNFa in the *Ccl19+* cluster^46^ and WNT signaling in the *Adamdec1+* cluster^22^, consistent with known biology (Extended Data Fig. 5g).

Differentially expressed genes (DEGs) in the *Pi16+* cluster relative to all clusters or the *Col15a1+* cluster alone suggested an identity similar to adventitial stromal cells (ASCs) and group 1 interstitial progenitors, including genes such as *Pi16, Dpp4* and *Ly6c^26,47,48^* (Extended Data Fig. 5h). ASCs and group 1 interstitial progenitor cells are cell states found across tissues^26,47,48^ that produce ECMs and have the capacity to acquire gene expression profiles consistent with development into more specialized fibroblast cell types^48,49^. The *Col15a1^+^* cluster exhibited an association with the basement membrane, such as *Col4a1, Hspg2* and *Col15a1,* a thin ECM-rich layer found within the parenchyma of tissues^50^ (Extended Data Fig. 5a and h). These data suggested that a primary role of these two cell states is to secrete and modify ECM in tissues across the body.

The ubiquity of the universal *Pi16+* and *Col15a1+* clusters across tissues and an elevated level of stemness-associated genes such as *Cd34* and *Ly6a* (encodes SCA-1), particularly in the *Pi16+* cluster, relative to other clusters (Extended Data Fig. 5i), led us to investigate the relationship between the fibroblast clusters. We applied the *Slingshot* algorithm^51^ to the steady-state fibroblast atlas and observed a trajectory that emerged from the *Pi16+* cluster, passed through the *Col15a1+* cluster, and ended at specialized clusters, such as the *Comp+, Ccl19+* or *Cxcl12+* via the *Adamdec1+* and *Coch+* clusters (Fig. 1g and Extended Data Fig. 5j). Collectively, our analysis showed that in steady-state mouse tissues, universal fibroblastic cell states (*Pi16+* and *Col15a1+*) and specialized fibroblasts exist and that these may be developmentally linked. These data also revealed that while bulk sequencing modalities suggested fibroblast identity is determined by tissue-type, analysis at the single-cell level showed instead that the fibroblast lineage is organized by functional specialization that is influenced, but not defined, by tissue.

### Cross-tissue transcriptional panorama of fibroblasts in perturbed mouse tissues

We next looked to determine how fibroblasts are impacted by infection^26^, injury^23,24,31,52^, cancer^12,21^, fibrosis^25,27,28^, metabolic changes^20,35^ and arthritis^7^ (Fig. 2a). Here, we integrated 13 publicly-available scRNA-seq datasets across 11 tissues implementing a strategy similar to our steady-state analysis to generate a perturbed-state fibroblast atlas (*n* = 67,868 cells) (Fig. 2b-c, Extended Data Fig. 6a-c, Supplementary Table 3). Using techniques similar to those applied to the steady-state fibroblast atlas, we identified and annotated 6 fibroblast clusters: *Pi16+, Col15a1+, Cxcl12+, Lrrc15+, Thbs4+, Cxcl5+* and a cluster of *Rgs5+* mural cells (Fig. 2b-c, Extended Data 6d, Supplementary Table 5). Similar to the steady-state fibroblasts, fibroblast clusters in the perturbed-state atlas displayed heterogeneous expression of common fibroblast genes (Fig. 2d). We observed that universal fibroblast subtypes and the *Cxcl12+* clusters in the perturbed-state atlas exhibited transcriptional similarities with the analogous clusters in the steady-state atlas (Fig. 2e). The *Cxcl12+* cluster has been previously reported to exhibit few transcriptional changes during inflammation so these cells were not examined further^21^, whereas a similar observation for the universal fibroblasts clusters has not been previously reported.

**Fig 2.**
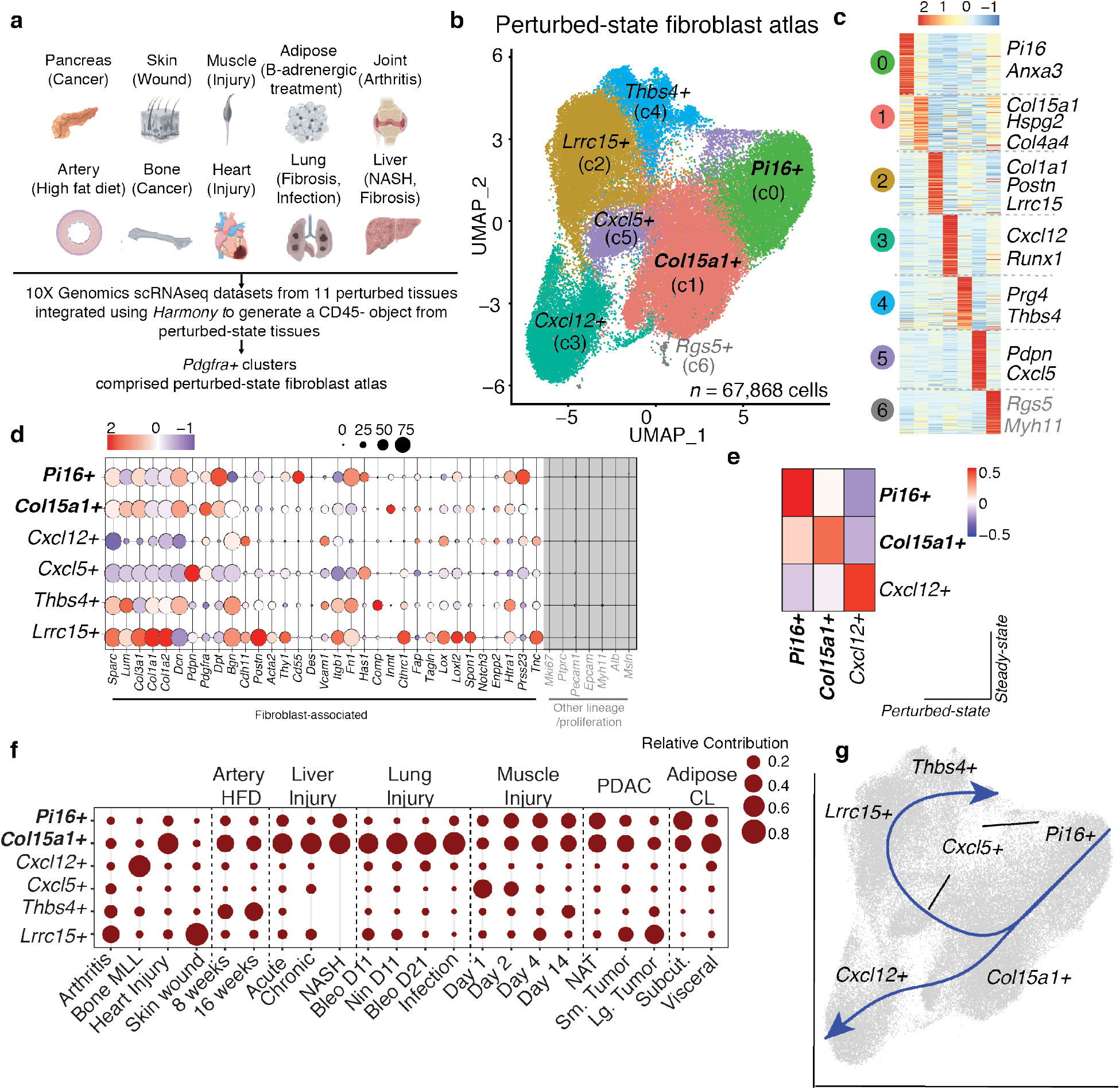
Cross-tissue transcriptional panorama of fibroblasts in perturbed mouse tissues. **a**. Schematic of datasets compiled for perturbation analysis from 11 perturbed mouse tissues: artery (high fat diet (8w and 16w), bone and bone marrow (mixed lineage leukemia), heart (injury), lung (fibrosis (bleomycin and bleomycin with nintedanib), parasite infection (*N. brasiliensis),* liver (NASH, Ccl4 treatment), pancreas (PDAC-GEMM model; Pdx1cre/+;LSL-KrasG12D/+;p16/p19flox/flox mice), skin (wound), muscle (notexin injury), visceral and subcutaneous adipose (beta-3 adrenergic treatment), joint (serum transfer induced arthritis). **b**. UMAP embedding of 67,868 cells in perturbed-state fibroblast atlas. Seven clusters identified through graph-based clustering are indicated by color. **c**. Heat map of the relative average expression of the most strongly enriched genes for each cluster identified by log-fold change of cells in one cluster compared to all other cells (z-score per row). **d**. Fibroblast- and other lineage-associated genes (in grey). The size of circles denotes the percentage of cells from each cluster, color encodes the average expression across all cells within a cluster. The color scale shows the expression level based on row z-score. **e**. Similarity between steady-state (x-axis) and perturbed state (y-axis) signature genes for *Pi16+, Col15a1+* and *Cxcl12+* clusters. Mean-centered values shown. **f**. Relative abundance of each tissue in steady-state UMAP clusters. The size of bubbles indicates the contribution of cells from each tissue to a cluster, columns without a bubble indicate lack of contribution (number of cells = 0) of that tissue to the corresponding cluster. Graph to be read column-wise. **g**. Pseudotime(s) visualized using principal curves representing two trajectories of fibroblast differentiation across steady-state fibroblast object. *Pi16+* cluster set as root fibroblast state.

In nearly all perturbed tissues and for all types of inflammation, some fibroblasts occupied the universal *Pi16+* and *Col15a1+* clusters (Fig. 2h and Extended Data Fig. 7a-v). The specialized clusters from the steady-state atlas, *Adamdec1+, Comp+, Ccl19+* and *Coch+,* did not have analogous clusters in this perturbed-state atlas, though this may be due to the composition of datasets within the perturbed-state analysis (Extended Data Fig. 6f). Conversely, projecting the *Cxcl5+, Thbs4+* and *Lrrc15+* clusters onto the steadystate map suggested that these clusters were observed only during inflammation (Extended Data Fig. 6g). Therefore, we surmised that *Cxcl5+, Thbs4+* and *Lrrc15+* clusters represented activated fibroblast states.

Fibroblasts from early muscle injury contributed a majority of cells to the *Cxcl5+* cluster. Cells within the *Thbs4+* cluster derived from arteries of mice on a high fat diet, muscle injury (*Day 14),* arthritis and large tumors from pancreatic ductal adenocarcinoma (PDAC). The *Lrrc15+* cluster was composed of cells from arthritis, skin wound, and small and large PDAC tumors (Fig. 2h and Extended Data Fig. 7a-v). In perturbed tissue samples, universal *Pi16+* fibroblasts maintained high levels of stemness-associated genes relative to other clusters, and *Col15a1+* fibroblasts displayed intermediate levels (Extended Data Fig. 6h). Consistent with this and in agreement with our observations in the steady-state, *Slingshot* trajectory inference again revealed a path from *Pi16+* cells through *Col15a1+* and then on to the specialized (*Cxcl12*+) or perturbation-specific, activated *Cxcl5+, Lrrc15+* and *Thbs4+* clusters (Fig. 2i and Extended Data 6i).

### Activated fibroblasts are comprised of *Cxcl5+* and *Thbs4+* fibroblasts and *Lrrc15+* myofibroblasts

We next examined the molecular signatures of activated *Cxcl5+, Thbs4+* and *Lrrc15+* clusters by comparing these to the universal *Pi16+* and *Col15a1+* populations. The *Cxcl5*+ cluster expressed lower levels of ECM-associated genes, collagens and matrix degrading proteins than the other clusters (Fig. 3a-e). Similar to the steady-state fibroblast atlas, we performed gene set enrichment analysis on the DEGs found between all clusters. This revealed that genes enriched in the *Cxcl5+* cluster were overrepresented in gene sets pertaining to inflammatory cytokines such as interleukins-1 and −6 (Extended Data Fig. 8a). Consistent with this, the *Cxcl5*+ cluster expressed chemokines such as *Ccl2* and *Ccl7^31,53^* that have been reported in acute muscle injury (Fig. 3d). Analysis of co-regulated genes suggested that this cluster was driven by TNFa and NFkB signaling (Fig. 3f). IL-1 cancer-associated fibroblast (CAF)^12,13^ and inflammatory fibroblasts in human RA^7^ have been previously described to secrete inflammatory factors that recruit immune cells, similar to the *Cxcl5*+ cluster. However, we observed that IL1 CAF were mostly restricted to the *Pi16+* universal fibroblast cluster, with some overlap into the *Cxcl5*+ cluster (Extended Data 8b). This suggests that in some cases cells described as inflammatory fibroblasts may represent *Pi16+* universal fibroblasts that have not undergone sufficient transcriptional change to constitute a new cell state, possibly due to localization distal to areas of active injury^8,16^.

**Fig 3.**
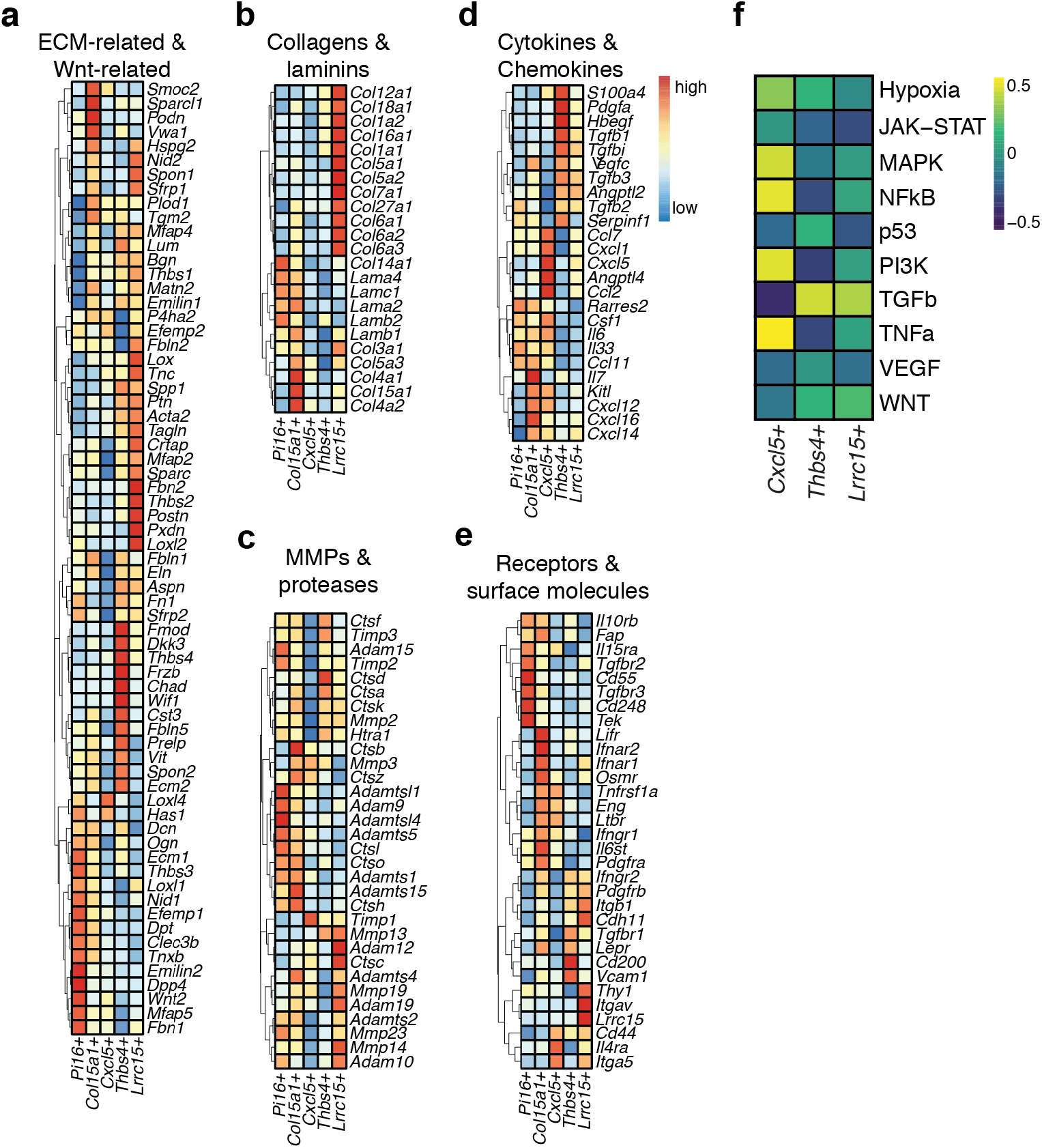
Activated fibroblasts are comprised of activated *Cxcl5+* and *Thbs4+* fibroblasts and *Lrrc15+* myofibroblasts. **a-e**. Heat maps showing average relative gene expression in *Pi16+, Col15a1+, Cxcl5+, Thbs4+* and *Lrrc15+* clusters (z-scored per row) in the following categories: **a**. ECM-related and Wnt-associated genes. **b**. Collagens and laminins. **c**. Matrix metalloproteases and cathespins. **d**. Cytokines and chemokines. **e**. Receptors and surface molecules. **f.** Expression of pathway-responsive genes in perturbed-state atlas clusters as assessed by PROGEN(y) analysis (z-scored per row).

Cells within the *Thbs4+* cluster exhibited elevated expression of *Prg4, Htra1* and *Hbegf,* suggesting that specifically lining fibroblasts from RA localized to the *Thbs4+* cluster^7,8,54^ (Fig. 3a,c,d). Enrichment of genes including *Cilp, Chad* and *Wif1* indicate that this cluster also contains cells approximating matrifibrocytes^55^ and Fibroblast-*Thbs4* and -*Clip*^56^, three cell states which have been identified as ECM producing and modifying cell states in the context of cardiac fibrosis.

The *Lrrc15+* cluster expressed uniquely high expression of an array of collagens, including fibrillar collagens, and *Acta2,* suggesting that at the transcriptional level cells in this cluster are myofibroblasts. Cells within the *Lrrc15+* myofibroblast cluster additionally differentially expressed glycoproteins such as *Cthrc1, Postn* and the metallopeptidase *Adam12,* relative to universal fibroblasts (Fig. 3a-c, e). This increase in fibrillar collagen expression is consistent with previous reports examining this cluster in PDAC^12^. As observed in the Thbs4+ cluster, pathway inference suggested that gene networks in both *Thbs4+* fibroblasts and *Lrrc15+* myofibroblasts are primarily driven by TGFb signaling, though we hypothesize that additional signals are required for development of these distinct fibroblast subtypes (Fig. 3f). Collectively, these data support the concept that universal fibroblasts may give rise to unique activation states (our perturbed-state atlas revealed three distinct activation states) which are not observed in steady-state conditions but only in contexts of injury and inflammation.

### Single-cell RNA-seq of human fibroblasts reveals universal and activated clusters across tissues and disease indications

We hypothesized that mice may exhibit some parity to humans in terms of steady- and perturbed-state fibroblast subtypes. To test this, we first performed scRNA-seq on tumor and normal adjacent tissue (NAT) samples from 3 patients with pancreatic cancer (*n* = 21,262 cells; Supplementary Table 6). We identified 22 clusters (Fig. 4a) which separated into immune cells, stromal cells, and epithelial cells using established markers^12^ (Extended Data Fig. 9a, Supplementary Table 7).

**Fig 4.**
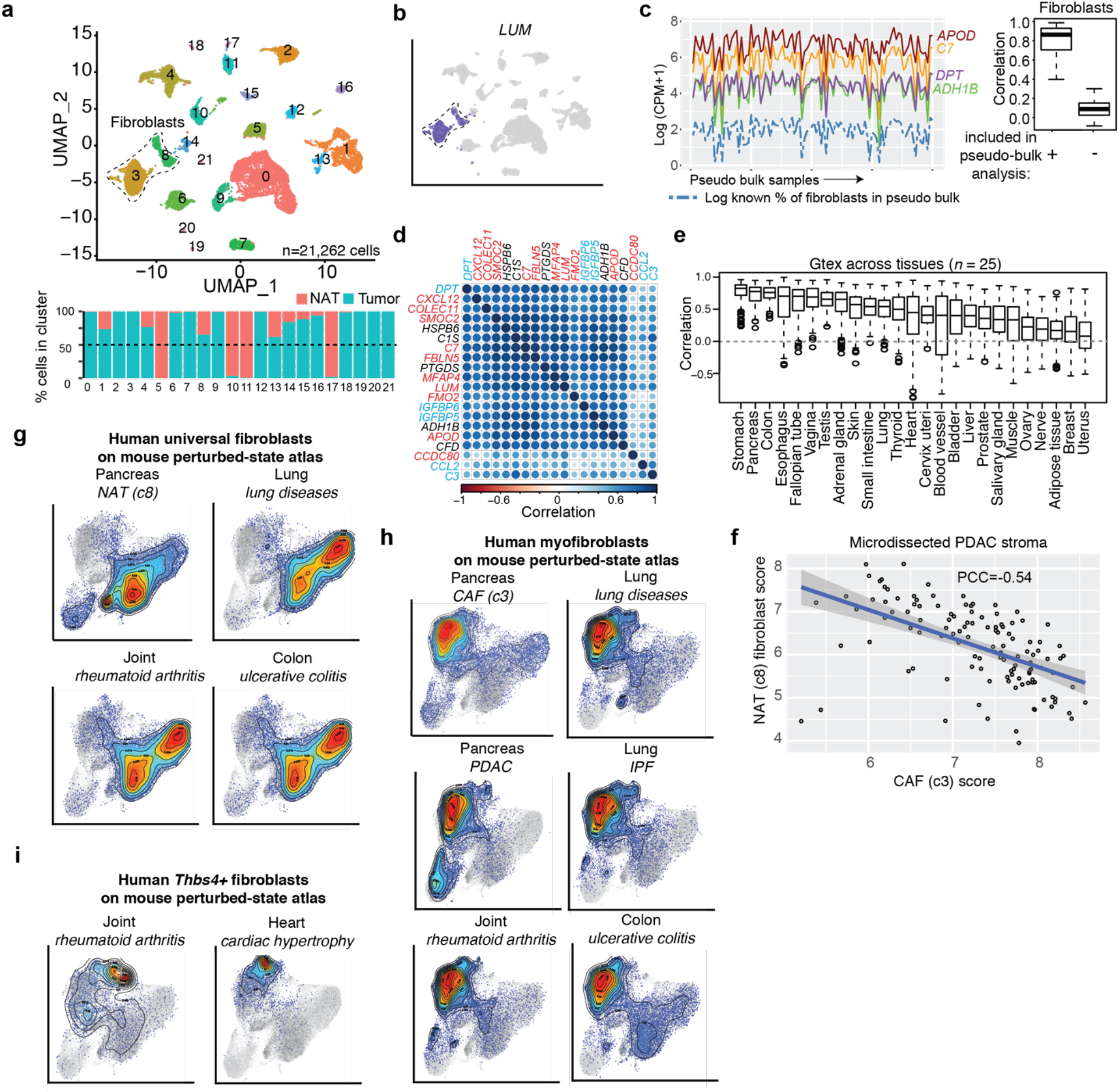
Human fibroblasts exhibit universal, specialized and activated cell types across tissues and disease indications. **a**. Top; UMAP embeddings of human pancreatic cancer tumor and normal adjacent tissue (n = 21,626 cells). Bottom: Percentage of cells in each cluster coming from Tumor or NAT. **b**. UMAP as in a, colored by expression of *Lum.* **c**. Left: Expression level of indicated marker genes (color, y-axis) across 100 pseudo-bulk samples (x-axis) generated from human PDAC scRNA-seq data. The known percentage of cells from cluster 8 in each pseudo-bulk is given by the dotted blue line. Right: Boxplots representing the distributions of pairwise correlation coefficients of the top 20 marker genes for cluster c8 in pseudo bulk samples containing (left) and not containing cells from c8 (right). **d**. Gene-by-gene correlation matrix of pairwise correlations in 205 normal pancreas bulk RNAseq samples from GTEx. Blue indicates *Pi16+* cluster signature gene, Red indicates *Col15a1+* signature gene. **e**. Co-expression as in d, results from the gene-by-gene correlation matrices are summarized as boxplots for each individual tissue from GTEx. **f**. Scatterplot visualizing the scores for a NAT (c3) fibroblasts expression geneset (y-axis) compared to a scores for CAF (c8) expression geneset (x-axis) in 122 bulk RNA-seq samples of microdissected PDAC tissue. Each dot represents a sample, the regression line is given in blue. **g-i**. UMAP representation of cells from the mouse perturbed-state atlas, each cell colored by their score for gene sets corresponding to: **g**. Universal fibroblasts from human PDAC (c8 NAT), RA (“THY1+CD34+”^7^), lung disease (“PLIN2+ fibroblast”^59^) and ulcerative colitis (“S1”^45^). **h**. *Lrrc15+* myofibroblasts from human pancreatic cancer (c3 CAF, “MyCAF”^13^), RA (“Human RA F2”^7^), lung disease (“Myofibroblast”^59^), IPF^60^ and ulcerative colitis (“S2”^45^). **i.** *Thbs4+* fibroblasts from human RA (“Lining”^54^) and cardiac hypertrophy^56^.

We classified clusters 3 (c3) and 8 (c8) as fibroblasts (Fig. 4b, Extended Data Fig. 9b). C3 was entirely composed of cells from tumor tissue and represented CAFs, while c8 was comprised of both tumor and NAT fibroblasts. Genes specific to *Pi16+* and *Col15a1+* universal fibroblasts in mouse (*DPT, C7, APOD;* Supplementary Table 4) were enriched in cluster 8 relative to cluster 3, suggesting that these might represent a less activated state (Supplementary Table 7). Currently, only limited fibroblast-enriched human single-cell datasets from a diversity of tissues are available, so atlases similar to what we show for mouse could not be constructed directly. Consistent with the observation of mouse universal genes enriched in cluster 8 compared to cluster 3, twelve of the 20 most upregulated genes in c8 fibroblasts compared to all cells in the dataset were significantly upregulated in the mouse steady-state *Pi16+ (DPT, IGFBP5, IGFBP6, C3*) or *Col15a1+ (CXCL12, SMOC2, C7, FBLN5, MFAP4, LUM, FMO2, APOD*) clusters. To test if cluster 8 cells from our PDAC single-cell data in fact represent universal fibroblasts in humans, we examined bulk RNA-seq data from a broad range of tissues (GTEx database (*n* = 5,961 samples).

To enable inference of universal fibroblast abundance in bulk RNA-seq data, we first showed that the 20 gene c8 signature derived from the single-cell data exhibited highly correlated expression pattern in human pseudo-bulk data. The expression levels of these genes tracked known abundance of cells from cluster 8 in the pseudo-bulk samples, and their strong-inter-gene correlation largely vanished when omitting cluster 8 from the pseudo-bulk samples (Fig. 4c). Thus, strength of pairwise correlation among these signature genes provides an indirect measurement of the abundance in a bulk RNA-seq sample. Confirming this result, the c8 gene signature was strongly correlated in normal human pancreas bulk RNAseq data from GTEx (Fig. 4d). Furthermore, they were undetectable in nearly all (96.3% ± 2.4%) of non-fibroblast cells in our human PDAC single-cell dataset. We carried out pairwise correlation analysis for these 20 markers in GTEx bulk RNA-seq data from 25 tissues chosen based on their high levels of fibroblast marker gene expression and found strong (PCC>0.5) co-expression in 12 human tissues (Fig. 4e and Extended Data Fig. 9c). Overall, these results suggest that human fibroblasts equivalents to both *Pi16+* and *Col15a1+* universal fibroblasts in mouse exist across tissues. These results align with observations in human adipose tissue^48^ and skin^57^ that have previously observed fibroblasts with transcriptional similarities to universal fibroblasts.

To characterize the cluster solely composed of CAFs (c3), we compared these cells to the c8 NAT enriched cluster (Fig. 4f). Many of the genes found in the mouse perturbed-state atlas *Lrrc15+* myofibroblast cluster were differentially expressed in c3 CAFs compared to c8 NAT cells, including *LRRC15, COL8A1, POSTN,* and *TAGLN* (Extended Data Fig. 9d). We next scored bulk RNAseq stromal samples from pancreatic cancer patients (*n* = 122)^58^ for the human universal fibroblast expression module (20 most enriched genes in c8 NAT vs c3 CAFs) as well as the activation program (20 most enriched genes in c3 CAFs vs c8 NAT). We found a strong negative correlation (PCC= – 0.54) between universal fibroblast score and the c8 CAF activation score, suggesting that acquisition of an activation program in human fibroblasts may be associated with loss of universal fibroblast gene expression (Fig. 4f).

To further support a hypothesized correspondence between clusters in mouse and human single-cell data, we next asked if the transcriptional fibroblast subtypes we observed in humans appeared to be present in mice. We found concordance between the human c8 NAT signature and the mouse universal *Col15a1+* fibroblast cluster (Fig. 4g). We also observed that the human c3 CAF gene signature was enriched in the murine *Lrrc15+* myofibroblast cluster (Fig. 4h). As the clusters in the mouse perturbed-state atlas are not restricted to tumor lesions but also are derived from multiple tissues and injuries, we scored cells in the perturbed-state atlas for fibroblast activation signatures that we derived from other human injuries. We found abundant evidence of universal fibroblasts in human joints derived from rheumatoid arthritis (RA) patients^54^, lung diseases^59^ and ulcerative colitis samples^45^, similar to what we observed in our human PDAC dataset (Fig. 4g). Fibroblasts from a previously published human pancreatic cancer dataset^13^, RA^7,10^, interstitial lung diseases^59^, IPF^60^ and ulcerative colitis^45^, exhibited transcriptomes that localized these cells principally to the mouse *Lrrc15+* myofibroblast cluster (Fig. 4h). We also found that human RA exhibited fibroblastic cells that corresponded to the mouse *Cxcl12+* and *Thbs4+* clusters^10,54^ (Fig. 4i and Extended Data Fig. 9e). Genes enriched in human cardiac hypertrophy were also enriched in the mouse *Thbs4+* cluster (Fig. 4i). As we observed in the mouse, cells considered to be inflammatory fibroblasts across human samples and indications only infrequently projected onto the mouse *Cxcl5+* cluster, similar to what we observed for IL-1 CAF (Extended Data Fig. 8b), most of these cells were restricted to the universal clusters^7,10,13,45,59^ (Extended Data Fig. 9f). Overall, these data point to similarities in patterns of fibroblast activation across human diseases that are also reflected in murine disease models.

## Discussion

Fibroblasts have emerged as nexus cells that augment the function and positioning of structural cells, immune cells and other cell types in steady-state conditions. Our findings, based on the analysis of 16 mouse tissues in the resting state indicated that *Pi16+* and *Col15a1+* universal fibroblasts reside in all organs. In contrast, some tissues – such as secondary lymphoid organs, intestine and bone – also possess specialized fibroblasts in the steady state. Fibroblasts are also drivers of pathology in a number of different disorders including arthritis, cancer and fibrotic diseases, and these cells can be activated by an array of cytokines and signaling pathways^61^. We found by investigating 11 tissues undergoing perturbation that universal fibroblasts can still be identified in perturbed tissues, along with three types of activated fibroblast cell types that emerge during inflammation. Lineage inference using *Slingshot* suggested a continuum of fibroblast differentiation in tissues, one in which the *Pi16+* cluster represents a root fibroblast state from which specialized or activated cells emerge after first transitioning through a *Col15a1+* intermediate state. Methods for inferring lineage from cross-sectional data can suggest likely patterns of differentiation, but true longitudinal studies will be required to confirm this hypothesis. This paradigm suggests a process by which fibroblasts populate tissues and respond to changes in the microenvironmental niches they inhabit. While the key niche factors required by most specialized and activated fibroblasts remain to be identified, these data suggest that these cells at once determine their microenvironment by promoting niches for other cell types and are yet acutely shaped by it.

Our data revealed concordance between fibroblast states in mice and in humans. First, we confirmed that a subset of fibroblasts in human patients has transcriptional similarities to the universal *Pi16+* and *Col15a1+* fibroblast cells we identified in mice, and that cells with similar transcriptional phenotypes are likely found within many of the 25 human tissues we examined. Second, examination of human datasets from indications such as pancreatic cancer, arthritis, interstitial lung diseases, IPF and ulcerative colitis revealed that clusters we identify in mice may have orthologs in humans, suggesting our mouse perturbed-state atlas represents a blueprint for fibroblast disease states in human diseases. Therefore, a deeper understanding of the development, evolution and molecular aspects of activated fibroblasts in mouse models may yield dividends in human medicine, though important differences no doubt exist as well.

The approach we took to understand fibroblasts across the body aimed to reveal broad similarities and essential differences between fibroblastic cell types. Our findings relied upon data we generated and upon deposited datasets contributed to public databases by other laboratories. Exposing the contours of fibroblast gene expression across tissues and activation states in this manner should be useful in achieving clarity on fibroblast subtyping and nomenclature. Open questions still remain, including elucidating the interplay and spatial dynamics between the fibroblast subtypes we identify, the existence of other potential subtypes not captured in these datasets, and the structural or immune cells that promote specialization in the steady state or activation during inflammation. It remains unclear as to why we observed two universal fibroblast states, though, as we observed evidence of these cell states across species, we speculate that these fibroblasts represent a necessary division of labor within the lineage. Overall, our data provide a framework for a deeper understanding of fibroblasts and may enable treatments focused on modulating these cells for clinical benefit.

## Supplemental Figures and Legends

**Extended Data Fig 1.**
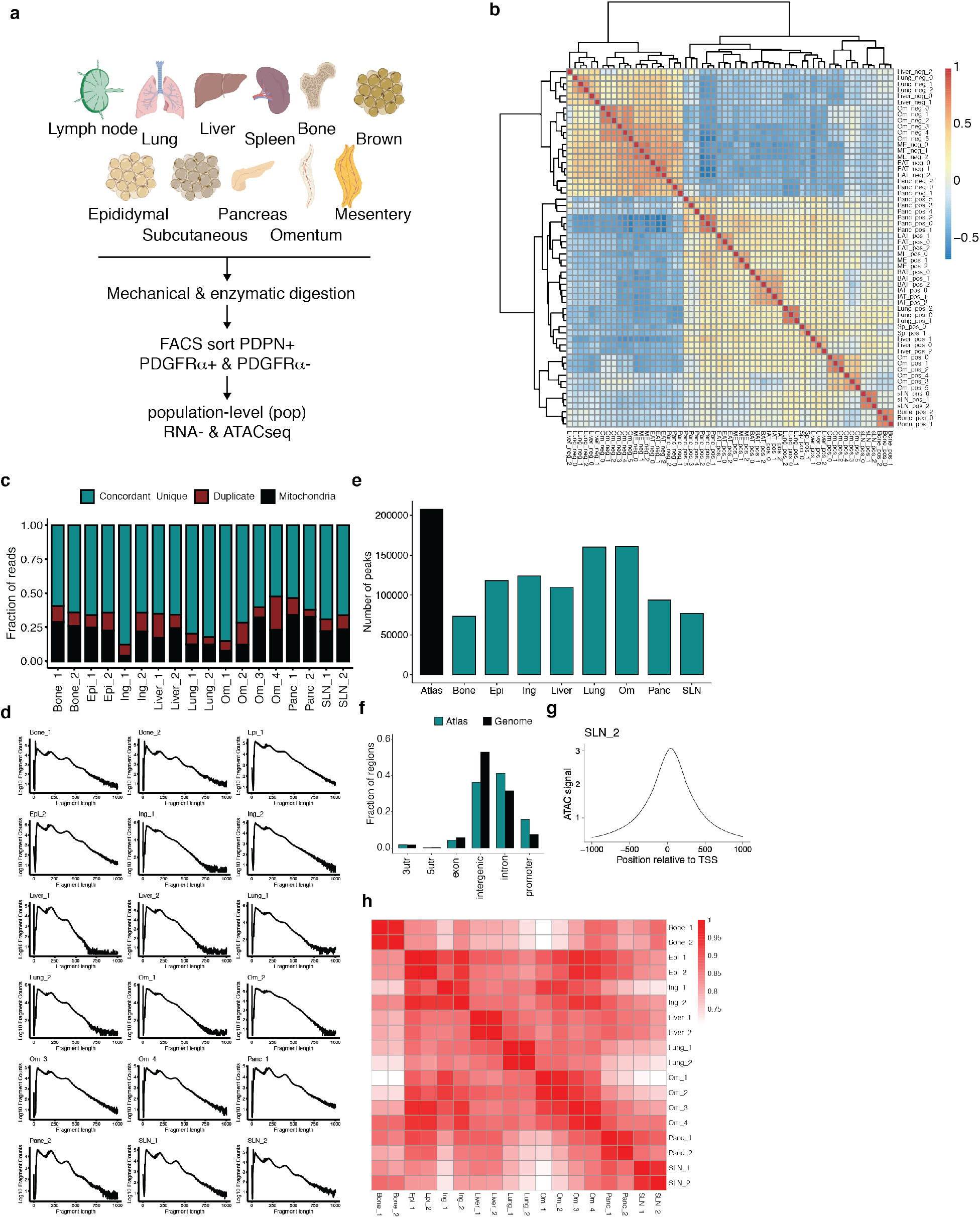
**a**. Diagram of tissues isolated for bulk RNAseq and ATACseq (adipose (brown, subcutaneous, epididymal), bone, liver, lung, lymph node, mesentery, omentum and pancreas) and experimental scheme. **b**. Correlation plot of bulk RNA sequencing samples based on top 1000 most differentially expressed genes. **c**. Fraction of ATACseq reads identified as PCR duplicate, mitochondrial DNA, or unique non-duplicate non-mitochondrial based on genomic mapping. **d**. Fragment lengths of ATACseq samples. **e**. Number of called peaks across ATACseq samples. Atlas describes the universe of all peaks. **f**. Distribution of ATAC-seq peaks across genomic regions. **g**. Aggregate signal around the transcription start site. sLN_2 samples shown, representative of all other samples. **h.** Heatmap of pairwise correlation coefficients of ATAC-seq samples.

**Extended Data Fig 2.**
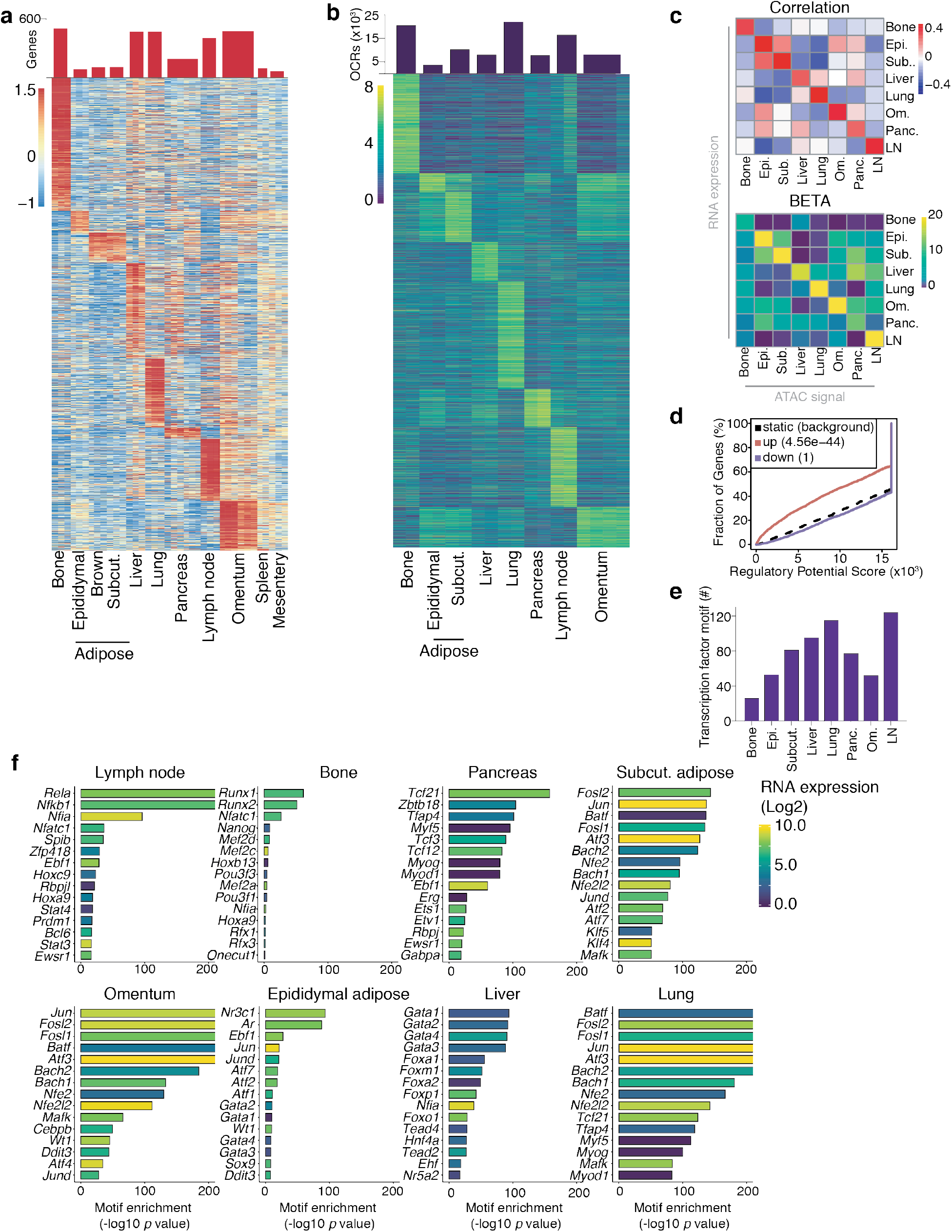
**a**. Heat map depicting enriched genes per tissue by bulk RNAseq. adj. *p* value < 0.05, Log2 FC > 2. Row z-scored. Top: barplot depicting number of signature genes per tissue. **b**. Heat map depicting regions of open chromatin per tissue by bulk ATACseq. adj. p value < 0.01, Log2 FC > 2. Row z-scored. Top: barplot depicting number of open chromatin regions per tissue. **c**. Correlation (top) and BETA analysis (bottom) of bulk RNAseq and ATACseq samples. **d**. BETA analysis of sLN evaluating enriched gene expression compared to enriched sLN OCRs. These data are representative of the rest of dataset. **e**. Number of transcription factor binding motifs in signature OCRs per tissue. **f.** Statistical inference of transcription factor motif enrichment in fibroblasts. Color of bar displays RNA expression of transcription factor.

**Extended Data Fig 3.**
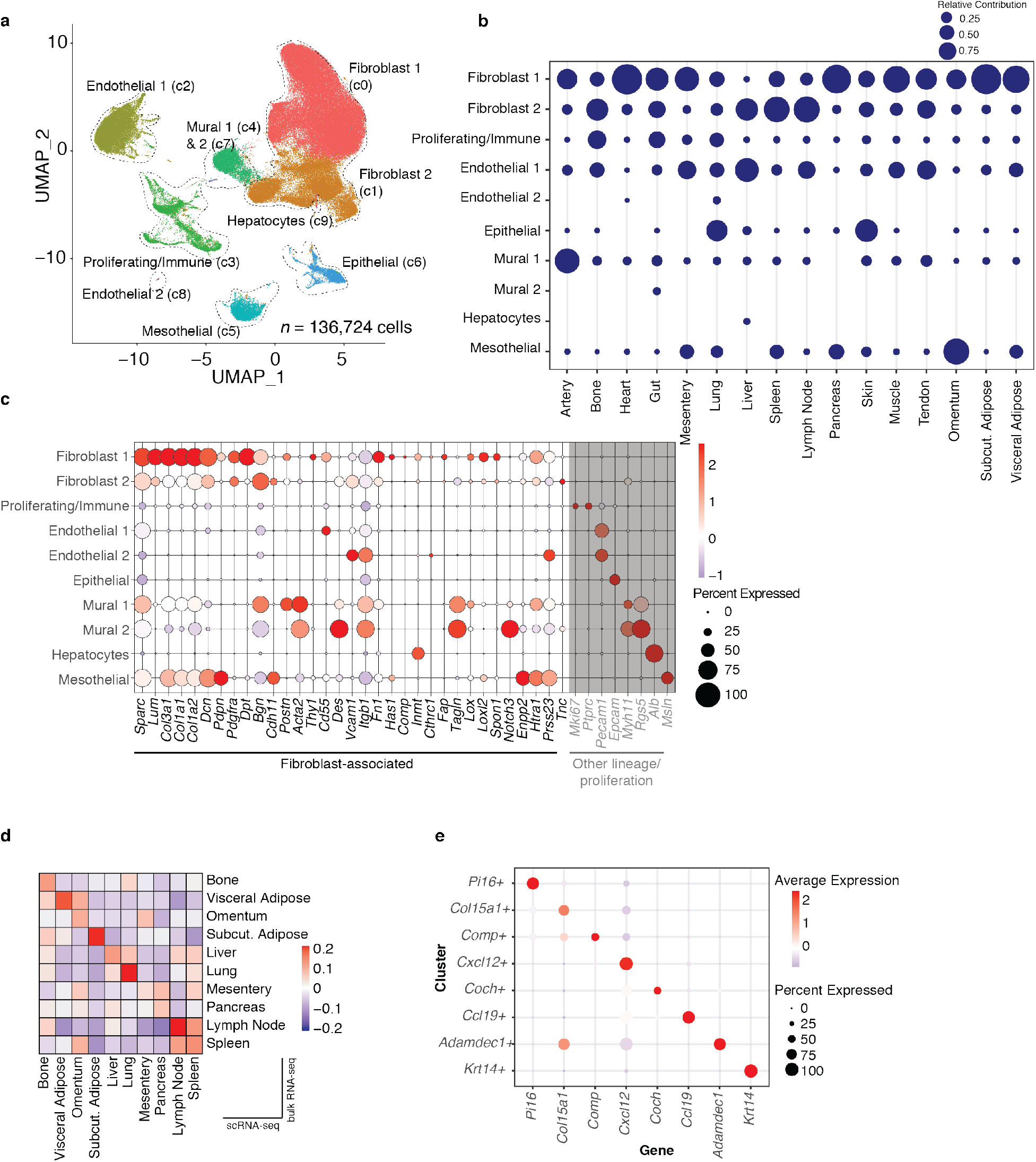
**a**. UMAP embedding of 136,724 single cells in steady-state CD45-atlas. Ten clusters identified through graph-based clustering are indicated by color. **b**. Relative abundance of each tissue in steady-state CD45-UMAP clusters. The size of bubbles indicates the contribution of cells from each tissue to a cluster, columns without a bubble indicate lack of contribution (number of cells = 0) of that tissue to the corresponding cluster. Graph to be read column-wise. **c**. Fibroblast- and other lineage-associated genes (in grey). The size of circles denotes the percentage of cells from each cluster, colour encodes the average expression across all cells within a cluster. The color scale shows the expression level based on row z-score. **d**. Average bulk tissue-specific fibroblast gene signature scores across tissues represented in the steady-state atlas. Mean-centered values shown. **e.** Expression of cluster hallmark genes in steady-state fibroblast atlas. The size of circles denotes the percentage of cells from each cluster, color encodes the average expression across all cells within a cluster. The color scale shows the expression level based on row z-score.

**Extended Data Fig 4.**
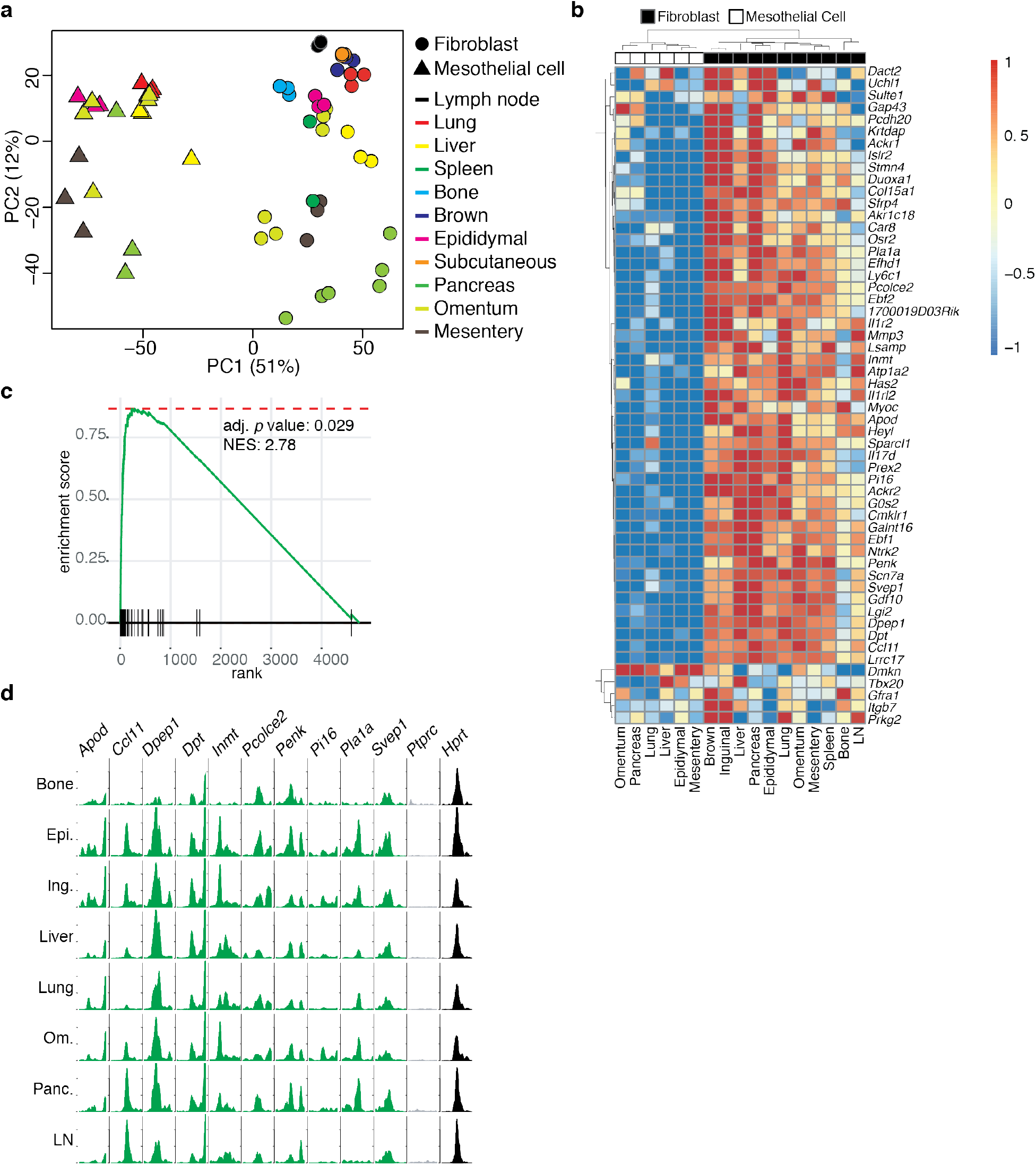
**a**. Principle component analysis of FACS-sorted bulk RNAseq of fibroblasts and mesothelial cells, calculated for the 1000 genes with the highest interquartile range. Circles represent fibroblasts and triangles are mesothelial cells. Each color denotes a different tissue. **b**. Heatmap depicting top 20 *Pi16+* and *Col15a1+* genes from steady-state fibroblast atlas in bulk RNAseq data. Rows are z-scored. **c**. Gene enrichment analysis of top genes from *Pi16*+ cluster and top 20 genes from *Col15a1+* cluster in loadings of PC1, which discriminates mesothelial cells and fibroblasts. **d.** ATACseq traces of select *Pi16+* and *Col15a1+* genes, *Ptprc* (encoding CD45) and Hprt at +/− 2kb of the TSS.

**Extended Data Fig 5.**
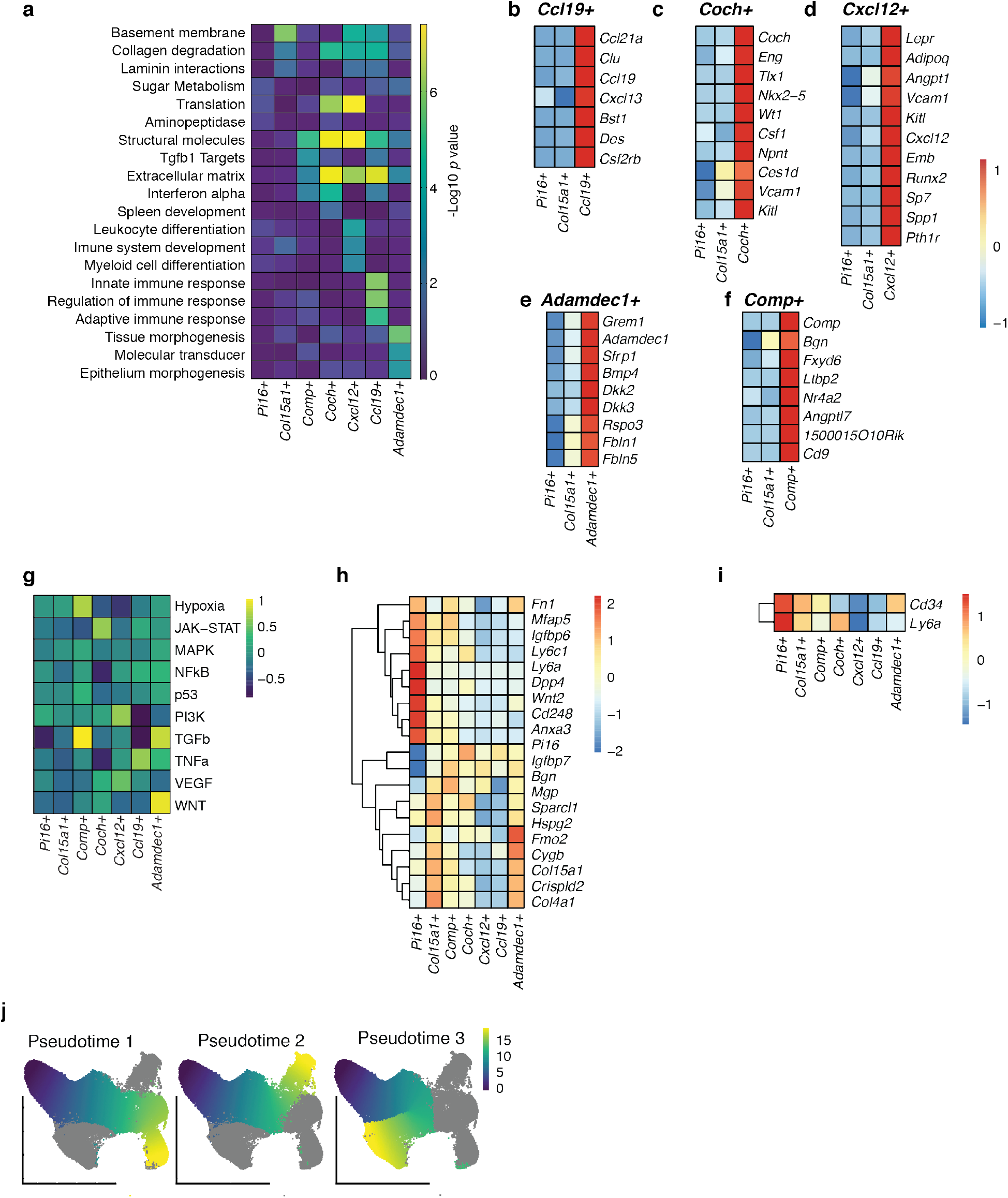
**a**. Gene set enrichment analysis of steady-state fibroblast clusters. Rows z-scored by negative log10 fold change. **b-f, h-j**. Heat maps of selected differentially expressed genes as noted below. All genes enriched at *p*<0.05, average log fold-change>0.25 unless otherwise noted. **b**. Select DEGs between *Pi16+* and *Col15a1+* clusters and *Ccl19+* cluster. **c**. Select DEGs between *Pi16+* and *Col15a1+* clusters and *Coch+* cluster. **d**. Select DEGs between *Pi16+* and *Col15a1+* clusters and *Cxcl12+* cluster. **e**. Select DEGs between *Pi16+* and *Col15a1+* clusters and *Adamdec1+* cluster. Rspo3 p = 1.04e-21, log fold change = 0.2495. **f**. Select DEGs between *Pi16+* and *Col15a1+* clusters and *Comp+* cluster. **g**. Expression of pathway-responsive genes in perturbed-state atlas clusters as assessed by PROGEN(y) analysis (z-scored per row). **h**. Select DEGs between *Pi16+* and *Col15a1+* clusters. **i**. *Ly6a* and *Cd34* expression in steady-state clusters. P< 0.05. **j.** UMAPs of three different lineages in the steady-state fibroblast atlas colored by pseudotime.

**Extended Data Fig 6.**
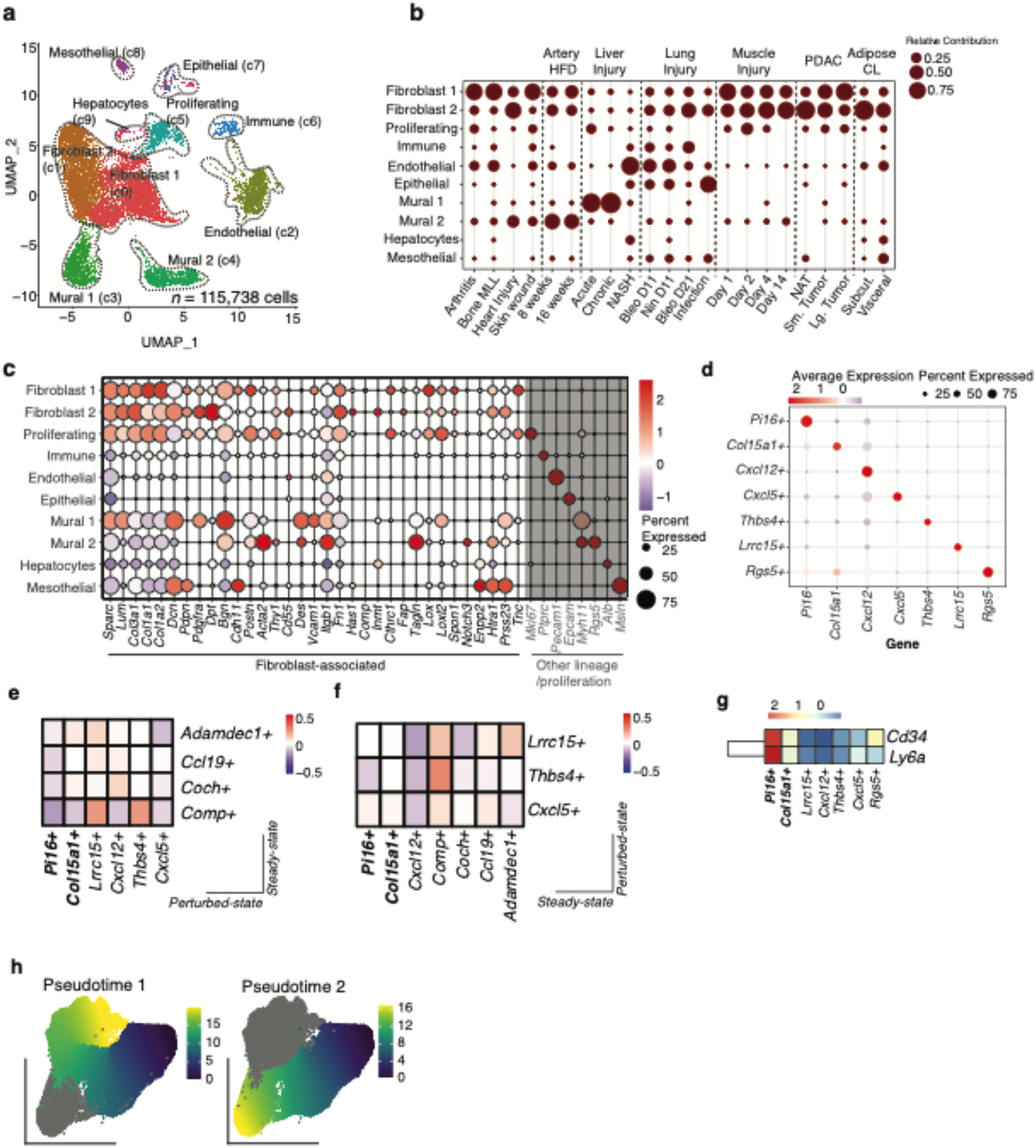
**a**. UMAP embedding of 115,738 cells in perturbed-state CD45-atlas. Ten clusters identified through graph-based clustering are indicated by color. **b**. Relative abundance of each tissue in perturbed-state CD45-UMAP clusters. The size of bubbles indicates the contribution of cells from each tissue to a cluster, columns without a bubble indicate lack of contribution (number of cells = 0) of that tissue to the corresponding cluster. Graph to be read column-wise. **c**. Fibroblast- and other lineage-associated genes (in grey). The size of circles denotes the percentage of cells from each cluster, color encodes the average expression across all cells within a cluster. The color scale shows the expression level based on row z-score. **d**. Expression of cluster hallmark genes in perturbed-state fibroblast atlas. The size of circles denotes the percentage of cells from each cluster, color encodes the average expression across all cells within a cluster. The color scale shows the expression level based on row z-score. **e**. Correlation between perturbed-state (x-axis) and steady-state (y-axis) signature genes between *Pi16+, Col15a1+, Lrrc15+, Cxcl12+, Thbs4+* and *Cxcl5+* clusters (perturbed-state) and *Adamdec1+, Ccl19+, Coch+* and *Comp+* clusters. Mean-centered values shown. **f**. Correlation between steady-state (x-axis) and perturbed-state (y-axis) signature genes between *Pi16+, Col15a1+, Cxcl12+, Comp+, Coch+, Ccl19+* and *Adamdec1+* clusters (steady-state) and Lrrc15+, *Thbs4+* and *Cxcl5+* clusters (perturbed-state). Mean-centered values shown. **g**. *Ly6a* and *Cd34* expression in perturbed-state clusters. P < 0.05. **h.** UMAPs of two different lineages in the perturbed-state fibroblast atlas colored by pseudotime.

**Extended Data Fig 7.**
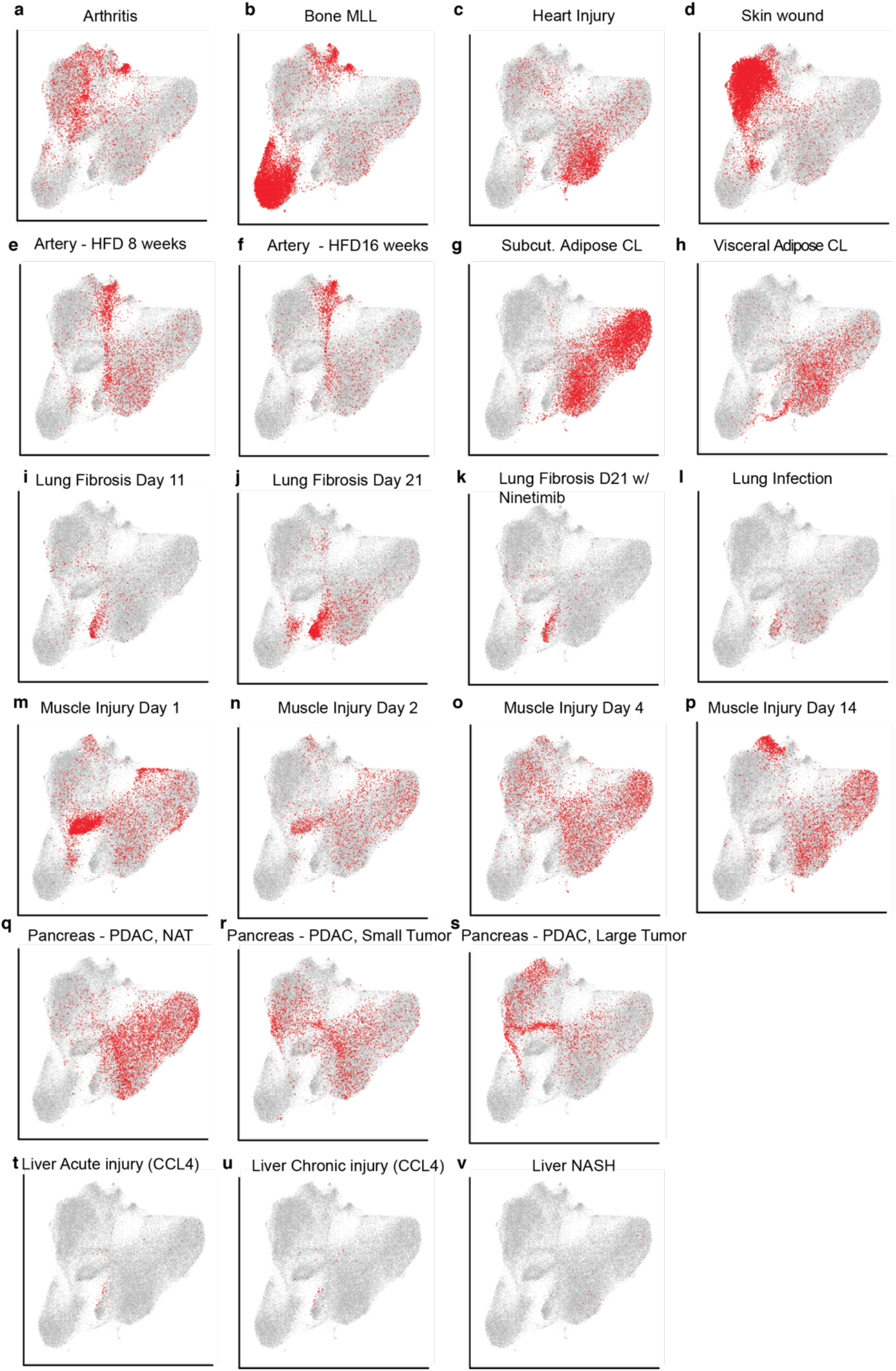
**a-v**. UMAP representation of the distribution of fibroblasts across tissues and perturbations.

**Extended Data Fig 8.**
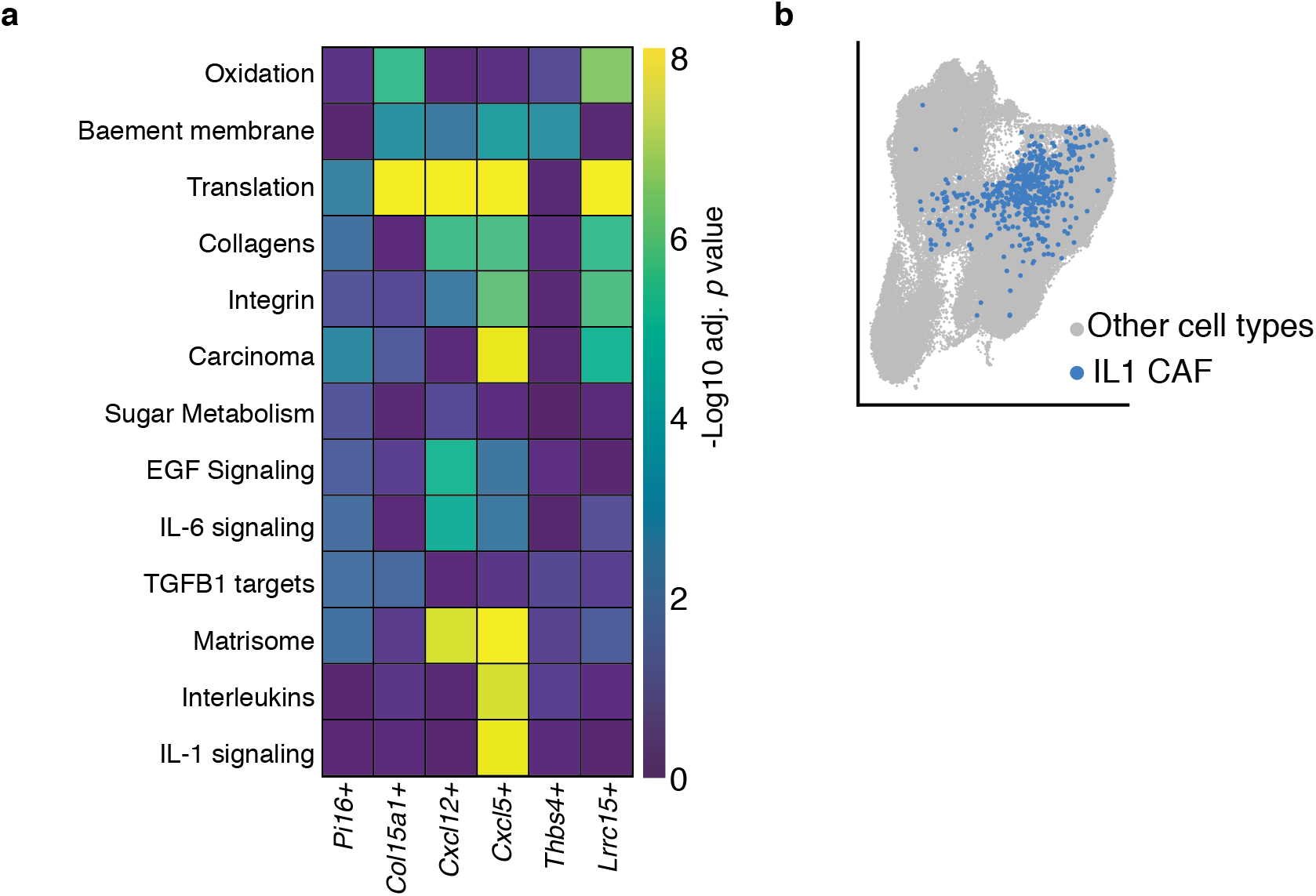
**a**. Gene set enrichment analysis of perturbed-state fibroblast clusters. Rows z-scored by negative log10 fold change. **b**. UMAP of IL-1 CAF cells derived from Dominguez and Muller et al., projected onto perturbed-state fibroblast atlas.

**Extended Data Fig 9.**
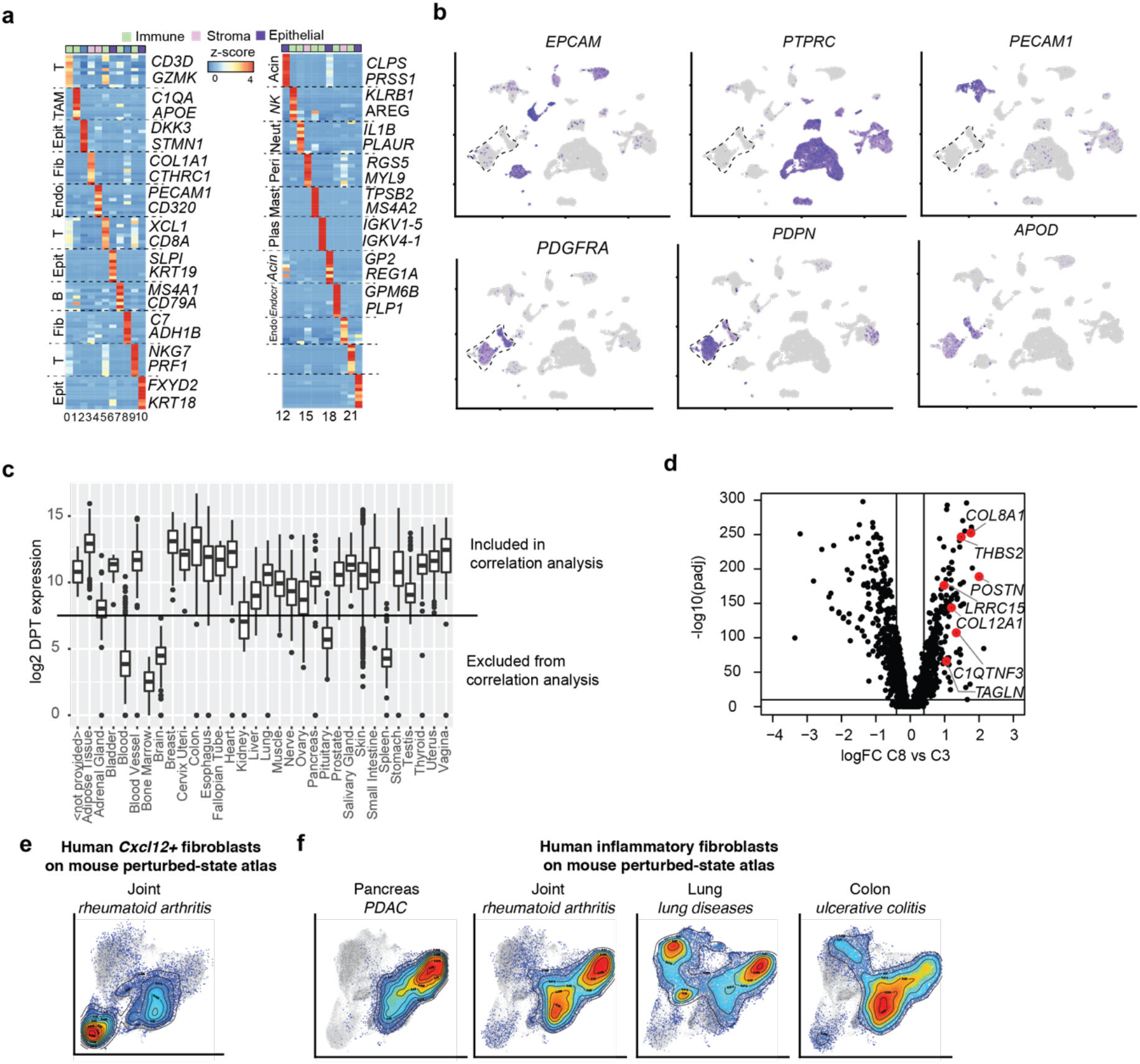
**a**. Relative average expression of top 10 marker genes (sorted by logFC) for each cluster in the PDAC single-cell dataset. Two representative genes highlighted per cluster. DEGs across clusters **b**. Representative featureplots **c**. Boxplots summarizing *DPT* expression distributions across tissues from the GTEx portal **d**. Volcano plot visualizing logFC (x-axis) and adjusted p-value (y-axis) comparing in fibroblasts from cluster 3 in panel A compared to fibroblasts from cluster 8. **e,f**. UMAP representation of cells from the mouse perturbed-state atlas, each cell colored by their score for gene sets corresponding to: e. *Cxcl12+* fibroblasts from RA (“HLA+ fibroblasts”^54^). **f.** Universal fibroblasts pancreatic cancer (“iCAF”^13^), RA (“Human RA F5”^7^), pulmonary fibrosis (“Has1 + fibroblast”^59^) and ulcerative colitis (“S4”^45^).

## Supplementary Tables

Table S1. (*Related to Extended Data Fig. 2*) Tissue-specific fibroblast and mesothelial cell genes by bulk RNAseq.

Table S2. (*Related to Extended Data Fig. 2*) Tissue-specific OCRs and TFs by bulk ATACseq

Table S3. (*Related to Fig. 1 and 2*) Tissue contributions to single cell RNAseq objects used in analyses of steady state and perturbed state murine tissues

Table S4. (*Related to Fig. 1*) DEGs for steady state fibroblast analyses

Table S5. (*Related to Fig. 2 and 3*) DEGs for perturbed state fibroblast analyses

Table S6. (*Related to Fig. 4*) Patient information

Table S7. (*Related to Fig. 4*) DEGs for human PDAC samples

Table S8. (*Related to Fig. 4*) Human gene sets projected onto mouse perturbed-state fibroblast atlas

## Methods

### Mice

Wildtype mice were obtained from Jackson Laboratory (JAX; colony 00064) and maintained at Genentech. Male mice, aged 6-12 weeks were used for all studies. All experiments were performed under protocols approved by the Institutional Animal Care and Use Committee at Genentech.

### Mouse tissue digestion and stromal cell isolation

Tissues were isolated and fibroblasts and mesothelial cells were isolated as previously described^17^. Briefly, tissues were obtained and minced, aside from the LN, omentum (neither minced) and bone (decapped, marrow removed and crushed). Next, tissues were placed in a 15mL conical tube with 5mL digestion media (RPMI + 2% FBS with 100mg/mL Dispase (Life Tech., cat. 17105041), 100-200mg/mL Collagenase P (Roche, cat. 11249002001), and 50mg/mL DNase I (Roche, cat. 10104159001)) and agitated. Tubes were placed in a 37C water bath for 15 mins, and 5mL fractions were removed and filtered (70uM) into RPMI supplemented with 2% FCS (VWR) three times. 200mg/mL Collagenase P was used to isolate cells from dense tissues such as the spleen, liver, inguinal adipose, brown adipose and pancreas and after a single cell suspension was obtained, the cells were layered on top of a 26% optiprep (Sigma, cat. D1556; diluted in PBS) gradient in 15mL conical tubes and spun at 1500 x g for 15 minutes with slow acceleration and the brake off. Cells in suspension were isolated with a transfer pipette. After digestion, the preparations were incubated with Ack for 2-5 minutes to remove red blood cells.

Cells were labeled with the following mAbs purchased from eBioscience, BioLegend, or BD Biosciences at 1:200 for 20-30 min, unless otherwise noted. Prior to cell surface staining with the following fluorescently labeled antibodies, cells were blocked with Fc block (2.4G2; 1:500-1:1000). Surface staining for experiments was performed as below, unless otherwise noted: CD45 (30-F11), EPCAM (G8.8), CD31 (390), PDGFRa (AP5), PDPN (8.1.1; 1:800). Live cells were identified by washing after Fc block and incubation with Fixable Viability Dye Violet (Invitrogen, cat. L34955, 1:1000) prior to surface staining or incubation with calcein blue (Invitrogen, cat. C1429, 1:1000) after surface staining. Data were acquired on a Fortessa, Symphony or LSRII (BD Biosciences) and analyzed using FlowJo (Tree Star) or cells were sorted on a Fusion or Aria (BD Biosciences).

### Human patient information

Pancreatic cancer sample collection was approved by the Ethics Committee of Beijing Cancer Hospital.

### Human tissue digestion and stromal cell isolation

Samples were obtained and sequenced by Analytical Biosciences. Single cells were dissociated from tumor and adjacent non-cancer tissues as described previously^62^. Briefly, tumors and adjacent non-cancer tissues were cut into approximately 1-2 mm^3^ pieces in the RPMI-1640 medium (Gibco), and enzymatically digested with gentleMACS (Miltenyi) for 60 min on a rotor at 37°C, according to manufacturer’s instruction. The dissociated cells were subsequently passed through a 100 μm SmartStrainer and centrifuged at 400 g for 5 min. After the supernatant was removed, the pelleted cells were suspended in red blood cell lysis buffer (TIANDZ) and incubated on ice for 1-2 min to lyse red blood cells. After washing twice with 1x PBS (Gibco), the cell pellets were resuspended in sorting buffer (PBS supplemented with 1% fetal bovine serum (FBS, Gibco)).

Single cell suspensions were stained with antibodies against CD45 and 7AAD for FACS sorting, performed on a BD Aria SORP instrument. Based on FACS analysis, single cells were sorted into 1.5 ml tubes (Eppendorf) and counted manually under the microscope. The concentration of single cell suspensions was adjusted to 500-1200 cells/ul. Cells were loaded between 7,000 and 15,000 cells/chip position using the 10x Chromium Single cell 5’ Library, Gel Bead & Multiplex Kit and Chip Kit (10x Genomics, V1.0 barcoding chemistry) according to the manufacturer’s instructions. All the subsequent steps were performed following the standard manufacturer’s protocols. Purified libraries were analyzed by an Illumina Hiseq X Ten sequencer with 150-bp paired-end reads.

### Mouse Bulk RNAseq analysis

For ex vivo bulk RNAseq, cells were isolated and stained as described above. Each tissue was represented by 2-3 individual replicates that were each derived by pooling tissues from 3-5 mice and FACS sorting cells directly into TRIZOL (Invitrogen, cat. 15596026). In some cases, lysed cells from at least three independent experiments were pooled for one replicate. In total, RNA was generated from an average of 35,195 +/7357 (SEM) fibroblasts and 17,318 +/− 7618 (SEM) mesothelial cells. RNA was isolated as described^63^ or at Expression Analysis, Inc.

Paired-end RNA-seq libraries were constructed from ≥747 pg of RNA using the SMART-Seq v4 ULTRA Low Input RNA Kit for Sequencing (Takara, cat. 634891) and NexteraXT kits (Illumina, cats. FC-131-1096 and FC-131-2001) for Low Input RNA Kits. Libraries were then sequenced on an Illumina HiSeq yielding, on average, 35 million read pairs (2×50bp) per sample. Reads were aligned to the GENCODE basic mouse transcriptome index (M14) and transcript levels quantified using salmon with parameters --type quasi -k 25. Subsequently, counts were transformed into gene-level counts in R using the *tximport* (https://bioconductor.org/packages/release/bioc/html/tximport.html) package. Differential expression analysis taking batches into account was carried out on the gene by sample count matrix with DESeq2^64^, using a design of ~0 + condition + batch having a coefficient for each level of condition. For principal component analysis, log transformed normalized counts (lengthScaledTPM) were batch corrected using Combat^65^ and PCA was performed in the space of variable genes (coefficient of variation >0.3). Gene set enrichment analysis (GSEA) using the fgsea method^66^ was performed on genes ranked by their principal component 1 loadings using the top 20 marker genes for *Pi16+* and *Col15a1+* clusters from the steady state fibroblast atlas.

### Mouse Bulk ATACseq analysis

For ex vivo bulk ATACseq, cells were isolated and stained as described above. Each tissue was represented by 2-4 individual replicates that were each derived by pooling tissues from 3-5 mice and FACS sorting fibroblasts. On average, 28455 cells (+/5325 (SEM)) were sorted per tissue. These cells were then frozen in Gibco Recovery Cell Culture Freezing Medium (ThermoFisher, cat. 12648010). The cells were then thawed in a 37°C water bath, pelleted, washed with cold PBS, and tagmented as previously described^67^, with some modifications^68^. Briefly, cell pellets were resuspended in lysis buffer, pelleted, and tagmented using the enzyme and buffer provided in the Nextera Library Prep Kit (Illumina, cat. FC-121-1031). Tagmented DNA was then purified using the MinElute PCR purification kit (Qiagen, cat. 28004), amplified with 10 cycles of PCR, and purified using Agencourt AMPure SPRI beads (Beckman Coulter, cat. A63882). Resulting material was quantified using the KAPA Library Quantification Kit for Illumina platforms (Roche, 07960255001), and sequenced with PE42 sequencing on the NextSeq 500 sequencer (Illumina), with 42-bp paired-end reads. Library preparation and sequencing was performed by ActiveMotif, Inc.

Reads were aligned to the GRCm38/mm10 build of the mouse genome using GSNAP^69^ with parameters -M 2 -n 10 -B 2 -i 1 --pairmax-dna=1000 --terminal-threshold=1000 --gmap-mode=none --clip-overlap. Read pairs aligning concordantly and uniquely to a single genomic location were retained for downstream analysis. PCR duplicates were removed using Picard MarkDuplicates (http://broadinstitute.github.io/picard/). Library depth corrected coverage bigwig files were obtained to visualize the regions of interest.

#### OCR identification

Open chromatin regions (OCRs) were identified as peaks on individual replicates and pooled samples combining the replicates of a given tissue using MACS2^70^, with parameters macs2 callpeak -f BAM --call-summits --nomodel --shift −95 --extsize 199 -- keep-dup all -p 0.1 --call-summits (these choices of the shift and extsize parameters correct for the +5/-4bp transposase insertion offset). The irreproducible discovery rate (IDR) pipeline^71^ was employed to assess peak concordance between the individual replicates of a given tissue, and these IDR estimates were subsequently appended to the associated pooled peaks. Robust peaks per tissue were defined as the pooled peaks that overlap at least 50% of a peak from at least 2 individual replicates and that pass an IDR threshold of 0.1. All robust peaks across all tissues were first centered on their summits (summit +/-199bp) and then those overlapping mitochondrial and noncanonical chromosomes were removed. Finally, all remaining peaks were merged to obtain the final set of all accessible regions (n = 207,803). Per sample, reads overlapping each region in the atlas were counted using the bedtools command multiBamCov^72^. To find tissuespecific OCRs, differential accessibility analysis was conducted on the count matrix using DESeq2^64^, where the accessibility (i.e., overlapping read count) of a given region in each tissue was compared against the count for that region in all other tissues. In this setting, the tissue-specific OCRs were defined according to the following criteria: log2 fold change >= 2, adjusted p-value <= 0.01. In addition, for each tissue a nondifferential/insignificant OCR set was defined according to: −0.585 <= log2 fold change <= 0.585, q-value > 0.05.

#### Motif enrichment analysis

For motif enrichment analysis, for each tissue-specific OCR set, an equally sized matched background set was selected based on region length and GC content from among the nondifferential/insignificant OCRs, using MatchIt^73^. AME^74^ from the MEME suite was used with default settings to assess the enrichment of a set of 321 PWMs from Homer (http://homer.ucsd.edu/homer/) in the tissue-specific OCR sets vs the background sets. Specifically, Fisher’s exact test was used to compare the number of matches to a given PWM in the specific set vs the background set, and assess statistical significance. Enriched PWMs were reported based on an adjusted p-value threshold of 0.05.

#### ATACseq and RNAseq concordance

To compute the correlation of log2 fold changes inferred from the ATAC-seq and RNA-seq differential analyses, the ATAC-seq final atlas peaks were assigned to the gene with the closest transcription start site (TSS), using Gencode mouse M14 annotations and a distance threshold of 50kb. Following the assignment, genes and atlas peaks with absolute log2 fold change >= 1 and q-value <= 0.05 in a given tissue were used in the correlation calculation.

An additional analysis to infer concordance between ATAC-seq and RNA-seq datasets was the BETA^75^ analysis, which takes a set of peaks (tissue-specific OCRs from ATAC-seq) and differential gene expression results from RNA-seq. In short, BETA calculates a regulatory potential score based on the number of peaks in a fixed window (100kb by default) around each gene TSS and ranks the genes based on this score. For each top gene set based on that rank, it calculates the percentage of the total up- and downregulated genes, as well as unregulated background genes to provide p-values for the overall up- or downregulation potential of the whole peak set. BETA was used with parameters -k BSF -g mm10 -n basic --df 0.1, for all pairwise tissue combinations, so for both matching and non-matching tissues.

### Mouse scRNA-seq meta-analysis

Integrated fibroblast atlases at steady- and perturbed-state were generated and analyzed using the following main steps (1) processing and filtering individual single-cell RNA-seq datasets from healthy and diseased tissues (2) integrating healthy and diseased datasets separately to generate steady- and perturbed-state atlases (3) clustering and annotation, (4) trajectory inference and, (5) gene set enrichment analysis for steady- and perturbed-state fibroblast atlases. The aforementioned steps are described in detail in the following sections.

#### (1) Processing and filtering individual single-cell RNA seq datasets

Single cell transcriptomics datasets, enriched in non-hematopoietic cells, generated using 10X Genomics and available as processed CellRanger files were collected from public repositories as well as from in-house lab data sets (Supplementary Table 3). For public datasets where, processed files were not made available, we analyzed raw data using cellranger count (CellRanger 2.1.0, 10x Genomics) using a custom reference package based on mouse reference genome GRCm38. A total of 23 single-cell RNA-seq data sets representing multiple tissues and perturbations were analyzed individually. In order to ensure comparability, for every individual dataset, we retained genes found in the Ensembl mouse (GRCm38) gene model, followed by implementing the Seurat single cell analysis pipeline (version 3.1.1)^76,77^ in R. Specifically, for each dataset we filtered low quality cells with < 500 measured genes and high percentage of mitochondrial contamination (> ~5-20 %, depending on the dataset). Post filtering, data in each cell was normalized to log(CPM/100+1), 2,000 most variable genes were identified, and the expression levels of these genes were scaled prior to performing PCA in the space of the most variable genes. Next, 20 principal components were used for graph-based clustering (resolution=0.1) and UMAP dimensionality reduction was computed. All steps were performed using functions implemented in the Seurat package (*NormalizeData, FindVariableFeatures, ScaleData, RunPCA, FindNeighbors, FindClusters, RunUMAP*) and default parameters, except where mentioned were used. Cell clusters marked by the canonical marker gene for immune cells, *Ptprc* (*Cd45*), were discarded. All individual datasets devoid of Cd45+ cells were next used for integration to create two main atlases: (1) a steady-state fibroblast atlas comprising data from healthy tissues, and (2) a perturbed-state fibroblast atlas comprising data from diseased and inflamed tissues.

#### (2) Dataset Integration for steady- and perturbed-state atlases

Prior to dataset integration, we imported aforementioned filtered, non-processed Seurat objects (not scaled) of healthy and diseased datasets and, determined a common gene space by retaining only those genes that were measured across all datasets (21,087 genes). Next, individual healthy and diseased Seurat objects were merged separately into two different steady- and perturbed-state objects, respectively. Each of these merged objects were normalized (function *NormalizeData,* method= “LogNormalize”, scale.factor = 10,000), scaled to regress out the stress gene signature (computed using Seurat’s *AddModuleScore*) of subpopulations affected by tissue dissociation methods^78^ prior to performing PCA for the most variable genes. These processed, merged objects were next used for batch effect correction and integration using Harmony^19^ (version 1.0). We adjusted the diversity clustering penalty parameter, theta to 1 in order to recapitulate all the cell populations from individual datasets without generating dataset specific clusters. We then provided the top 20 harmony dimensions as an input for UMAP and visualized the first two UMAP dimensions at a clustering resolution of 0.2. Next, we identified distinct cell types using canonical marker genes such as *Sparc, Col3a1, Dcn* (fibroblasts), *Epcam* (epithelial cells)*, Alb* (hepatocytes), *Pecam1* (endothelial cells), *Msln* (mesothelial cells), *Rgs5* (mural cells – pericytes), *Myh11* (mural cells – smooth muscle cells), *Top2a* and *Mki67* (proliferating cells), and *Cd24a* (remnant immune cell populations) (Supplemental Figures 3 and 6). The computational pipeline for integration was iterated twice to generate fibroblast atlases.

Specifically, at each of the following steps non-relevant/ unwanted cell types were filtered followed by batch-effect correction to generate fibroblast-specific atlas for steady- and perturbed state: (1) a CD45- steady- and perturbed state atlas comprising 136,724 and 115, 738 cells respectively (2) a non-endothelial stromal (NES) cell steady- and perturbed state atlas comprising 106,855 and 92,503 cells respectively (data not shown) from the CD45- atlas after discarding endothelial, epithelial and proliferating cells, and (3) a fibroblast steady- and perturbed state atlas comprising 86,379 and 67,868 cells respectively, after discarding other stromal cells including mesothelial cells, pericytes, smooth muscle cells from the NES atlas.

#### (3) Clustering and annotation of steady- and perturbed-state fibroblast atlas

For each fibroblast map, resolution of 0.2 was chosen, differential gene expression computed using the Seurat function *FindAllMarkers* using a Wilcoxon Rank Sum Test and corrected for multiple testing using the Benjamini-Hochberg method.

Gene expression scores were computed using Seurat’s *AddModuleScore* function, visualized using *VlnPlot* or *DotPlot.* To determine markers for specialized or activated clusters relative to universal fibroblasts we used the Seurat function *FindMarkers* with default parameters. Next, we scored bulk tissue specific signatures in the steady-state atlas, computed average scores per tissue signature across tissues represented in the steady-state atlas and visualized using the ComplexHeatmap function *Heatmap.* Steady and perturbed-state cluster analogs were determined by scoring steady-state cluster markers in the perturbed-state atlas, followed by visualizing the median score per cluster using the ComplexHeatmap function *Heatmap.* The reverse (perturbed-state cluster signatures in steady-state atlas) was also performed and visualized as a heatmap.

To infer the activity of signaling pathways that govern different fibroblastic cells at steady state and after perturbation, we implemented the Bioconductor package PROGENy^79^. For both fibroblast atlases, we implemented the same strategy. First, we down-sampled each atlas using the Seurat function *subset* with parameters ‘ *WhichCells(object, downsample = 1000, seed = 1)’* followed by implementing the function *progeny* with default parameters *‘scale = TRUE, organism = “Mouse”, top=100, perm=1, return.assay=TRUE’*. We then summarized the progeny scores by cell population and visualized them as a heatmap using the function *pheatmap.*

#### (4) Pseudotime reconstruction and trajectory inference

Single cell pseudotime trajectories for both steady and perturbed state maps were computed using the algorithm slingshot (version 1.2.0) which enables computation of lineage structures in a low dimensional space^51^. Specifically, slingshot was implemented in the analysis pipeline after dimensionality reduction and clustering of the integrated object. Pre-computed cell embeddings and clusters from the Seurat pipeline served as an input to the function *slingshot* (reducedDim = ‘UMAP’, clusterlabels = object$ RNA_snn_res.0.2, start.clus = “Pi16”, extendi =‘n’, stretch=0), with Pi16 cluster set as the root of the trajectory. The start cluster was chosen based on prior biological knowledge and the expression of genes such as *Cd34* and *Ly6a,* known markers of progenitor-like cells. The wrapper function slingshot then performed lineage inference by treating clusters as nodes and constructing a minimum spanning tree (MST) between the nodes. Next, lineages or trajectories were defined by ordering clusters via tracing paths through the MST. Finally, individual pseudotime(s) were visualized using principal curves.

#### (5) Gene Set Enrichment analysis

Functional enrichment analysis was performed using the Bioconductor package fgsea (version 1.10.1)^66^. Ranked gene expression markers computed using the Seurat function *FindMarkers* (logFC = 0) were used as an input for the *fgsea* wrapper function. The *fgseaMultilevel* function was provided with the ranked gene list along with a set of genesets constituting pathways from the msigdbr (species= “Mus Musculus”, category = “C2”), pathways with a gene set size greater than 500 and fewer than 15 were excluded from the analysis. Pathway data for all the clusters was collated, biologically meaningful and significant (*p* value < 0.05) pathways were selected and visualized as a heatmap (Prism, Graphpad).

### Bioinformatics data processing of human data

Raw sequencing data was transformed into FASTQ format with CellRanger’s (v2.1) *mkfastq* command, mapped to the human genome (GRCh38), and quantified with CellRanger *count* using default parameters. Quantified UMI count matrices from each patient were merged in R and analyzed with the Seurat package (v 3.1.4). First, cells with <500 measured genes, or <2,700 UMIs, or >10% mitochondrial counts were removed from the dataset. In the resulting filtered dataset, data in each cell was normalized to log(CPM/100+1), the 2,000 most variable genes were identified, and the expression levels of these genes was scaled prior to PCA in the space of the most variable genes. Subsequently, 30 principal components were used for graph-based clustering (resolution=0.1) and UMAP dimensionality reduction. All steps were performed with the methods implemented in the Seurat package (*NormalizeData, FindVariableFeatures, ScaleData, RunPCA, FindNeighbors, FindClusters, RunUMAP*) and default parameters, except for parameters mentioned above. Markers for each cluster were identified using the *FindAllMarkers* function, limiting the maximum number of cells per cluster to 1,000 for runtime improvement. Genes differentially expressed between clusters 3 and 8 were detected using the *FindMarkers* function and default parameters. To map human expression signatures onto the mouse perturbed state map, human gene symbols were translated to their mouse orthologs and an enrichment score for the gene signature was calculated using Seurat’s *addModuleScore* function. Gene sets were identified within referenced papers (Supplementary Table 8).

Pseudo-bulk samples for co-expression analysis were generated from the human single-cell dataset using the following strategy: We randomly sampled 10% of cells from the PDAC single-cell dataset and pooled their reads into a bulk profile, which was subsequently normalized to log2(CPM). Using this strategy, we generated 100 bulk RNA-seq profiles with known proportions of cells from individual single-cell clusters. This allowed us to compare the expression of individual cluster 8 marker genes across pseudo-bulk samples a) pairwise between genes and b) to the known cell type proportion of cluster 8 in the pseudo-bulks. Next, we generated similar bulk samples, but this time excluding cells from cluster 8 in the sampling process. On these samples we again calculated gene-by-gene correlation coefficients for C8 marker genes and compared the distributions of pairwise correlation coefficients to the distributions in the pseudo bulk containing cells from cluster 8.

GTEx bulk RNA-seq data of normal tissues was obtained as batch-corrected, log normalized counts from the UCSC Xenabrowser^80^. Pairwise correlations were visualized with the *corrplot* (https://cran.r-project.org/web/packages/corrplot/) package. For crosstissue correlation analyses, only tissues with a median *DPT* expression > 7.5 were considered. In this analysis, the top 20 marker genes for cluster 8 of the single-cell dataset ordered by logFC, which were found in less than 15% of other cells, were used. For deconvolution of microdissected PDAC stromal samples, raw expression counts per sample (n = 122) were downloaded from GEO (GSE93326). Data was normalized to Log2(CPM+1). Scores for c8 and c3-derived expression signatures (described above) in these bulk samples were calculated based on the average expression of the 20 most upregulated genes from the respective single-cell cluster (ordered by logFC, only genes expressed in at most 30% of other cells were considered).

#### Pseudo-bulk analytical strategy

We first generated 100 pseudo bulk RNA-seq profiles from our single-cell dataset with varying numbers of cells from individual single-cell clusters (Fig. 4c, left). We observed that the expression of marker genes for fibroblast cluster 8 co-varied depending on the number of cells from cluster 8 in the bulks. As a consequence, their expression profiles were strongly correlated, but only if cells from cluster 8 were added to the pseudo bulk. Leaving cells from cluster 8 out resulted in an extensive drop of gene-wise correlations close to 0. Therefore, co-expression of a single-cell derived marker gene set can be used to infer presence or absence of a particular cell population in bulk RNA-seq.

#### Projection of human gene sets onto mouse perturbed-state atlas

Gene expression signatures from human single-cell RNA-seq datasets (Supplementary table 7) corresponding to different fibroblast types were scored on the perturbed state atlas using the Seurat function *AddModuleScore*. The density of cells with the highest activation score (25th percentile for all clusters except in the Thbs4 cluster [where we visualized the top 5th percentile]) was visualized using the function *LSD::Heatscatter*.

### Data availability

Raw and processed RNAseq and ATACseq files are available from ArrayExpress repository under the accession numbers EMATB-xxxxand EMTAB-xxxx. Supplementary Table 3 lists the studies and GSE numbers used to generate the CD45-maps and fibroblast atlases. These integrated single-cell RNAseq objects used for analysis are provided in an online resource that can be accessed via: [link made available upon publication]

Human pancreatic cancer single-cell data is available in EGA under accession: EGASxxxxxxxx.

### Code availability

Code will be made available upon publication.

## Acknowledgements

We acknowledge members of the Turley laboratory and individuals in the bioinformatics department for their useful discussions. We thank facility staff at Genentech for vivarium maintenance and core facility assistance. We express gratitude to Turley laboratory members and AMB for their careful review of this manuscript. This work was supported by Genentech.

## Author contributions

Conceptualization: MBB, RNP, SM, RB and SJT. Methodology: MBB, RNP, SM, RB and SJT. Software, Formal Analysis and Data Curation: RNP, SM, MBB and ACK.

Investigation: RNP, SM, MBB and ACK. Writing: MBB, RNP, SM, RB and SJT. Visualization: MBB, RNP, SM. Supervision: SM, RB and SJT.

## Competing interests

All authors except for R. Bourgon are employees of Genentech. R. Bourgon is an employee of Freenome.

## Materials and Correspondence

Further information and requests for resources should be directed to Shannon Turley.

## Notes

### Competing Interest Statement

The authors have declared no competing interest.

